# Combined JAK2 and HSP90 inhibition produces synergistic anti-tumour activity across preclinical glioblastoma models

**DOI:** 10.64898/2026.11.18.767500

**Authors:** Muhammad Furqan, Richard J. R. Elliott, Alexander E.P. Loftus, Wai Kit Chan, Roza Masalmeh, Martin Lee, Morwenna T. Muir, Fraser Laing, John C. Dawson, Peter W.K. Nagle, Roderick N. Carter, Bhuvaneish T Selvaraj, Paul Baxter, Alison F. Munro, Camilla Drake, Vanessa Smer-Barreto, Gillian M. Morrison, Steven M. Pollard, John O. Mason, Valerie G. Brunton, Margaret C. Frame, Neil O. Carragher

## Abstract

**Background:** Glioblastoma (GBM) remains difficult to treat because of tumour heterogeneity and the limited activity of single-target therapies. Our previous phenotypic drug screen identified JAK2/STAT3 signalling and HSP90-dependent protein homeostasis as pharmacological vulnerabilities. We investigated whether combined JAK2 and HSP90 inhibition could produce broader anti-tumour activity across GBM models.

**Methods:** Multiple JAK2 and HSP90 inhibitors were tested alone and in combination across 2- and 3-dimensional patient-derived GBM stem cell models. Drug interactions were quantified using the zero-interaction potency (ZIP) synergy model. Mechanistic studies included RNA sequencing, flow cytometry, western blotting, immunofluorescence and FUCCI live-cell imaging. Activity was also assessed in cerebral organoid–GBM co-cultures, non-malignant human neural and glial cells and orthotopic GBM xenografts.

**Results:** Combined JAK2 and HSP90 inhibition produced synergistic interactions in 12 of the 13 GBM cell lines tested, using multiple structurally distinct inhibitor pairs. Gandotinib and NMS-E973 were selected for further mechanistic studies, where the combination reduced spheroid growth and increased caspase-3 activation and PARP cleavage. Combination treatment was associated with G2/M accumulation, altered mitotic regulation, abnormal chromosome and spindle organisation, and increased γH2AX signalling. Anti-tumour combination activity was maintained in cerebral organoid co-cultures, where tumour burden was reduced as assessed by imaging, histological and luciferase-based measurements, while the drug combination had relatively limited effects on non-malignant human neural and glial cell types tested over the same concentration range. *In vivo*, combined Pacritinib and NMS-E973 treatment significantly reduced tumour burden in orthotopic G7 xenografts compared with vehicle or either single agent.

**Conclusions:** JAK2/HSP90 co-inhibition produces reproducible anti-tumour activity across heterogeneous GBM models and disrupts cell-cycle and mitotic progression. These findings support further evaluation of this combination strategy in GBM.

## Introduction

Glioblastoma (GBM) is the most aggressive primary brain tumour in adults and remains difficult to treat^1^. The current standard of care includes maximal safe surgical resection followed by radiotherapy and temozolomide (TMZ).^1,2^ Despite this, recurrence is common and long-term survival remains poor.^1,2^ Several features of GBM contribute to treatment failure, including extensive infiltration into the surrounding brain tissue, limited drug delivery across the blood brain barrier, significant inter and intra tumour heterogeneity, and ability of GBM cells to rapidly change their cellular state and signalling in response to treatment.^1,3,4^ Together, these features allow different tumour cell populations to survive therapeutic pressure and thus make it difficult to achieve durable responses using therapies directed against a single molecular target.^4^ Patient derived GBM stem cell lines recapitulate the heterogeneity of primary tumour samples and are valuable tools for identifying vulnerabilities shared across different tumour backgrounds.^5^ In our previous phenotypic drug screening study, we tested 3,866 compounds across six patient-derived GBM lines and identified several pharmacologically tractable vulnerabilities, including compounds targeting JAK2/STAT3 signalling and proteins involved in protein folding and homeostasis, such as HSP90 and HSF.^5^ Since GBM cells depend on multiple interconnected signalling pathways for survival, we reasoned that combining vulnerabilities identified independently in this screen could produce stronger and broader activity across heterogenous GBM models than targeting either pathway alone.

The JAK2/STAT3 pathway has previously been linked to GBM cell survival and maintenance of stem like tumour cells.^6–8^ Activation of JAK2 leads to phosphorylation of STAT3 at Tyr705, allowing STAT3 to regulate transcriptional programmes involved in proliferation, survival and cell-state maintenance.^7^ Increased JAK2/STAT3 activity has been reported in GBM tissue, patient-derived tumour stem cells and orthotopic tumour models.^6^ Genetic or pharmacological inhibition of this pathway can reduce GBM cell survival, although responses differ between models.^6^ Pacritinib, a JAK2-targeting inhibitor, has shown anti-proliferative activity against patient-derived GBM cells and measurable brain exposure *in vivo*, although monotherapy did not significantly improve survival in an orthotopic GBM model.^9^ More broadly, JAK/STAT signalling operates within a highly interconnected network of oncogenic and microenvironmental pathways in GBM, where signalling redundancy and tumour heterogeneity can limit the effectiveness of targeting a single pathway.^7^ Reviews of JAK/STAT-directed therapies in GBM have therefore highlighted rational combination strategies as an important approach for improving therapeutic responses and limiting adaptive resistance.^7^ Together, these findings support JAK2/STAT3 as a relevant therapeutic vulnerability in GBM, while also providing a rationale for combining JAK2 inhibition with complementary treatments.

HSP90 represents a different but potentially complementary vulnerability.^10^ HSP90 is an ATP-dependent molecular chaperone required for stability and function of many proteins involved in cancer signalling^10,11^. Its client proteins include components of receptor tyrosine kinase, PI3K–AKT, RAF– MEK–ERK and JAK–STAT signalling pathways, allowing HSP90 inhibition to affect several oncogenic pathways at the same time^10,11^. In GBM models, HSP90 inhibition has been shown to reduce tumour cell growth and survival and to interfere with pathways involved in treatment resistance, and HSP90 inhibitors were also among the more active compound classes in our previous phenotypic screen^5,12^. Clinical development of earlier HSP90 inhibitors was limited by toxicity and variable efficacy; however, next generation HSP90 inhibitors have shown improved clinical tolerability^10^. Pimitespib (TAS-116), an orally available HSP90α/β inhibitor, showed a manageable safety profile and improved progression free survival in a phase III trial in advanced gastrointestinal stromal tumours, leading to its approval in Japan^13,14^. NMS-E973 is another HSP90 inhibitor with favourable pharmacokinetic properties in preclinical models.^11^ Previous studies have shown sustained tumour exposure and measurable levels of NMS-E973 in brain tissue, together with activity in an intracranial tumour model.^11^ These properties make NMS-E973 useful for investigating HSP90 inhibition in the context of GBM.

There is also a biological link between JAK2 and HSP90 that provides a rationale for combining inhibitors of these two targets.^15^ JAK2 has been identified as an HSP90 client protein, and inhibition of HSP90 can destabilise JAK2 and reduce downstream JAK–STAT signalling.^15^ In JAK2-dependent haematological malignancies, HSP90 inhibition has enhanced the effects of JAK2 inhibitors and can retain activity in models with reduced sensitivity to JAK2-targeted treatment^16^. These observations suggest that HSP90 inhibition may complement direct JAK2 inhibition by both weakening JAK2/STAT3 signalling and disrupting additional survival pathways maintained by HSP90^15,16^. Combination of JAK2 and HSP90 inhibitors has demonstrated promising *in vitro* and *in vivo* activity in chronic myeloid leukaemia models,^17^ but we are not aware of any studies investigating this interaction in heterogenous GBM models. Here, we investigated combined pharmacological inhibition of JAK2 and HSP90 across a panel of patient-derived GBM stem cell 2D growth assays using several inhibitors of both target classes. We then examined the effects of the combination on 3D spheroid growth, apoptotic cell death and cell-cycle regulation, including associated mitotic and γH2AX responses. We then extended our evaluation to more complex models, including human cerebral organoid–GBM co-cultures, and compared drug responses with non-malignant human neural and glial cell types. Finally, we evaluated the combination in an orthotopic GBM xenograft model.

## Results

### JAK2 and HSP90 inhibitor combinations show synergistic activity across GBM models

In our previous phenotypic drug-screening study, we identified several druggable vulnerabilities across a panel of patient-derived GBM stem cell lines, including compounds targeting JAK2/STAT3 signalling and HSP90 dependent protein homeostasis.^5^ We therefore asked whether combining inhibitors of these two target classes could improve anti-tumour activity across different GBM models. We first tested three JAK2 inhibitors (Gandotinib, Pacritinib and Fedratinib) and four HSP90 inhibitors (NMS-E973, Pimitespib, Ganetespib and AUY922) as single agents across a panel of patient-derived GBM stem cell lines from the Glioma Cellular Genetics Resource (GCGR). Sensitivity varied between GBM lines, although HSP90 inhibitors were generally more potent than JAK2 inhibitors (Fig. 1A). Among the HSP90 inhibitors, Ganetespib and AUY922 generally showed greater potency than NMS-E973 and Pimitespib across the GCGR panel. A similar pattern was observed in G7 3D spheroids, where Pimitespib showed the lowest pIC₅₀ among the HSP90 inhibitors tested (Fig. 1B). We used G7 spheroids as a complementary 3D model to extend our findings from the 2D screening assays and to support subsequent studies in cerebral organoid co-cultures and orthotopic xenografts. The corresponding dose-response curves are shown in Supplementary Fig. 1A,B. Gandotinib reduced STAT3 Y705 phosphorylation, while NMS-E973 altered AKT and ERK signalling (Supplementary Fig. 1C,D). These changes were consistent with modulation of known pathways affected by the two inhibitor classes. ^11,18^

**Figure 1.**
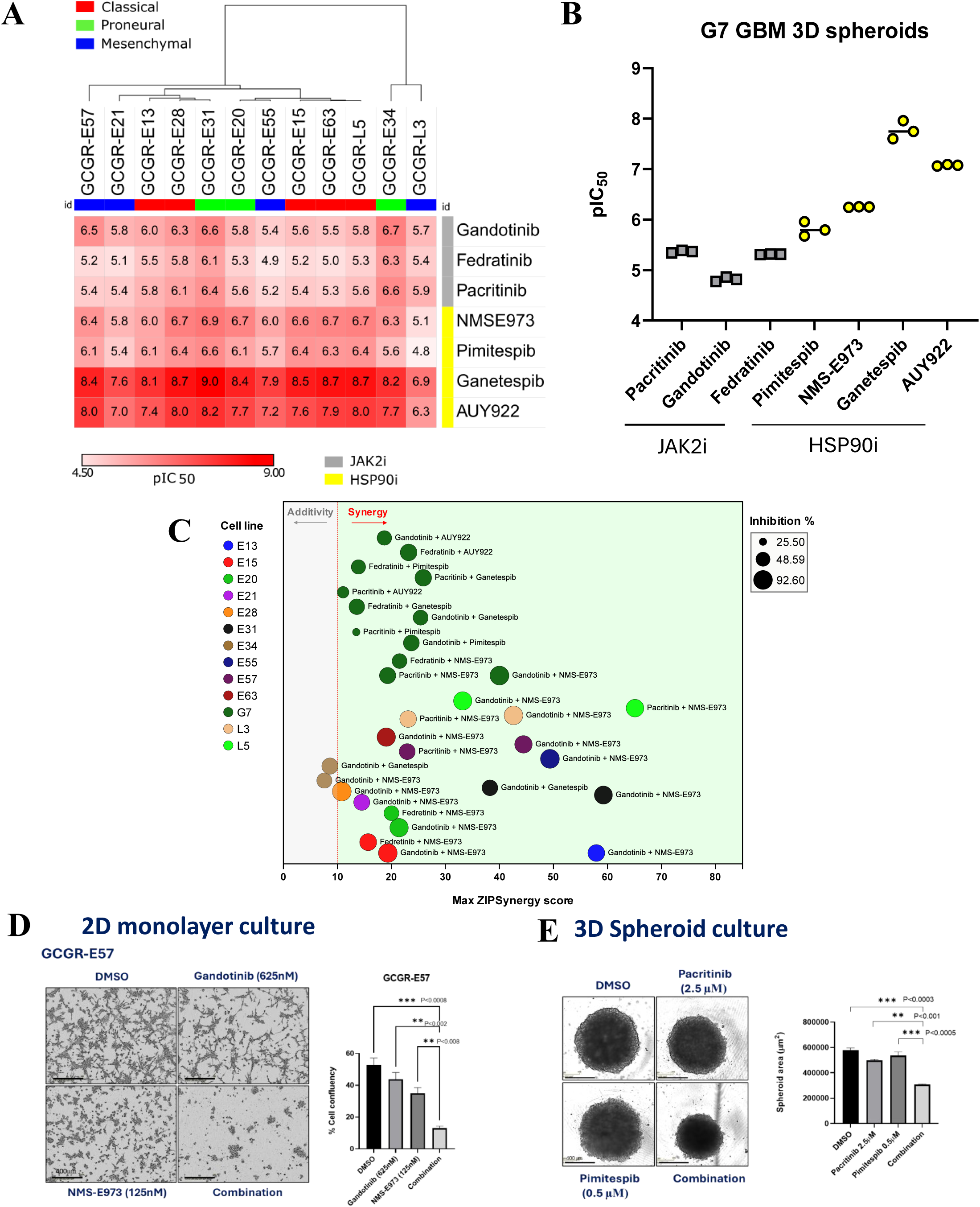
JAK2 and HSP90 inhibitor combinations show synergistic activity across GBM models. **(A)** Heatmap of pIC₅₀ values for JAK2 and HSP90 inhibitors across patient-derived GCGR GBM cell lines cultured under 2D conditions, determined using nuclei-based viability measurements. Heatmap colours represent pIC₅₀ values, ranging from 4.5 (white) to 9.0 (red), with higher values indicating greater compound potency. Compounds are grouped according to their target and cell lines are hierarchically clustered according to sensitivity. **(B)** pIC₅₀ values for the indicated JAK2 (grey squares) and HSP90 (yellow circles) inhibitors in G7 3D spheroids, determined using the PrestoBlue cell viability assay following 72 h treatment. Each dot represents an independent experiment (n = 3), and horizontal lines indicate the mean pIC₅₀ **(C)** Maximum ZIP synergy scores for JAK2/HSP90 inhibitor combinations across 13 GBM cell lines. Bubble size indicates percentage inhibition at the concentration inhibitor pair showing maximum ZIP score. Dotted line indicates a synergy threshold of ZIP score 10. **(D)** Representative images of GCGR-E57 cells treated with indicated compounds. Right: quantification of relative cell confluency after 72 hours of treatment. **(E)** Representative images of G7 3D spheroids treated with indicated compounds. Right: quantification of spheroid area (µm²) after 5 days of treatment. Data are shown as mean ± SD (n = 2 technical replicates per condition in each of three independent experiments). Scale bar, 400

We next tested JAK2 and HSP90 inhibitors in pairwise dose-ratio matrix combinations and measured drug interactions using the ZIP model.^19^ Synergy was observed across several GBM lines and with different JAK2-HSP90 inhibitor pairs, although the level of synergy varied between cell lines (Fig. 1C).

Synergistic interactions were observed in 12 of the 13 GBM cell lines tested, with maximum ZIP scores above 10 for at least one JAK2/HSP90 inhibitor pair (Fig. 1C). Only GCGR-E34 remained within the additive range. Gandotinib and NMS-E973 combination showed consistent activity across several cell lines and was therefore selected for further studies. Synergy was also observed in G7 spheroids with Pacritinib and Pimitespib (Supplementary Fig. 1E), as well as Gandotinib and NMS-E973 (Supplementary Fig. 1F), further suggesting that the effect was not limited to one specific compound pair. We next examined pathway modulation at Gandotinib/NMS-E973 concentrations associated with increasing levels of synergy in G7 cells (Supplementary Fig. 1G). Combination pairs with ZIP scores of 24, 38 and 54 were associated with progressive inhibition of STAT3 signalling as the concentration of Gandotinib increased. Interestingly, NMS-E973 treatment alone increased phospho-STAT3 Y705. To determine how early this response occurred, we performed a short time course analysis and found that increased STAT3 Y705 phosphorylation was detectable from 2 h following NMS-E973 treatment and remained obvious at 3 h (Supplementary Fig. 1H). This early increase in STAT3 phosphorylation is consistent with an adaptive signalling response to HSP90 inhibition.^20,21^ Importantly, the increase in phospho-STAT3 following NMS-E973 treatment was reduced when NMS-E973 was combined with Gandotinib while phospho-ERK was reduced following treatment with either Gandotinib or NMS-E973, without a clear additional reduction in the combination. The total STAT3, AKT and ERK levels remained relatively stable. These results show that synergistic drug concentrations were associated with simultaneous modulation of known JAK2/STAT3 and HSP90-dependent signalling pathways.

The effect of the combination was also observed across diverse image-based phenotypic assays. In GCGR-E57 cells, Gandotinib and NMS-E973 combination reduced cell confluency greater than either drug alone (Fig. 1D). Similarly, combined Pacritinib and Pimitespib treatment reduced G7 spheroid area compared with either single agent (Fig. 1E). In a complementary Matrigel-embedded G7 spheroid invasion assay, combination treatment also reduced spheroid outgrowth into the surrounding matrix (Supplementary Fig. 2A,B). Together, these results show reproducible synergistic activity between JAK2 and HSP90 inhibitors across multiple human GBM stem cell models.

### Combined JAK2 and HSP90 inhibition promotes apoptotic cell death in GBM models

We next asked whether the reduction in GBM spheroid growth following combined JAK2 and HSP90 inhibition was associated with apoptotic cell death. In G7 spheroids, caspase-3 activity remained relatively low following DMSO or single agent treatment, whereas combination caused a progressive increase in caspase-3 associated fluorescence over time (Fig. 2A). At 48 h, caspase-3-associated fluorescence was higher in combination-treated spheroids than in the other treatment groups. Representative images from the same live-cell imaging experiment showed stronger red caspase-associated fluorescence in combination-treated G7 spheroids compared with DMSO or either single agent (Fig. 2B). To examine whether the apoptotic response extended to other patient-derived GBM lines, including GCGR-E34 which showed an additive rather than synergistic response in the drug combination screen, we assessed PARP cleavage in GCGR-E15 and GCGR-E34 cells. In both cell lines, combination treatment increased cleaved PARP after 48 h compared with control and single agent treatment (Fig. 2C). We then measured caspase-3 activation in a wider panel of patient derived GBM lines. Although the timing and level of the response differed between cell lines, combination treatment generally caused higher caspase-3 activation than either drug alone in E13, E28, E21, E57 and E31 cells (Supplementary Fig. 3). Together, these results show that combined JAK2 and HSP90 inhibition promotes an apoptotic response across multiple GBM stem cell lines and growth contexts.

**Figure 2.**
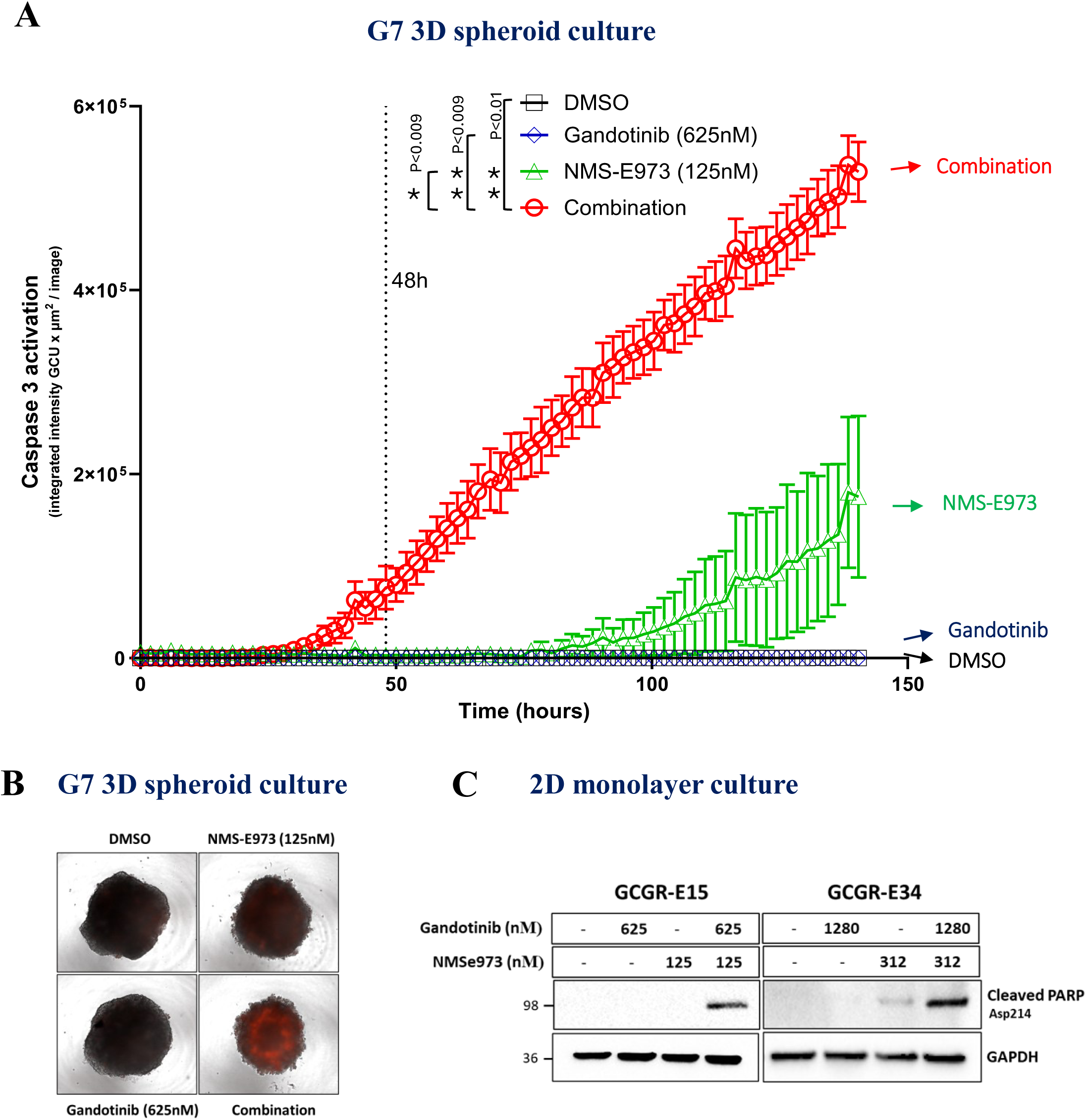
Combined JAK2 and HSP90 inhibition induces apoptotic cell death in GBM cells. **(A)** Live-cell quantification of caspase-3 activation in G7 3D spheroids treated with DMSO, Gandotinib (625 nM), NMS-E973 (125 nM), or the combination. Caspase-3 activity was monitored using BioTrac 530 Red Caspase-3 Dye and quantified as integrated red fluorescence intensity over time. Statistical significance was assessed at 48 h timepoint using one-way ANOVA with Dunnett’s multiple comparisons test; P values are indicated on the graph. Data are shown as mean ± SD, n = 3 technical replicates per condition in each of three independent experiments. **(B)** Representative brightfield and red fluorescence overlay images from the live-cell caspase-3 assay shown in (A), illustrating caspase-3-associated fluorescence in G7 spheroids following the indicated treatments at 140 hours **(C)** Western blot analysis of cleaved PARP in GCGR-E15 and GCGR-E34 cells following 48 h treatment with indicated drugs and control.

### Combined JAK2 and HSP90 inhibition alters cell-cycle regulation and promotes G2/M accumulation in G7 spheroids

To investigate the cellular response to combined JAK2 and HSP90 inhibition, we first performed RNA-seq analysis of G7 spheroids after 24 h treatment. Principal component analysis showed separation of combination-treated samples from DMSO and single-agent conditions, indicating a distinct transcriptional response (Supplementary Fig. 4A). To infer potentially differentially-regulated biological processes altered by treatment, we performed gene set variation analysis (GSVA)^22^ on a curated collection of 50 gene sets (MSigDB)^23^. This showed clear changes in cell-cycle-associated pathways, including increased G2/M checkpoint, E2F target and MYC target signatures in Gandotinib-treated samples, with a further increase following combination treatment (Fig. 3A; Supplementary Fig. 4B,C). Interestingly, this was not accompanied by a similar increase in the mitotic spindle signature. Mitotic spindle gene-set activity was increased following Gandotinib treatment alone but not in the combination-treated samples (Supplementary Fig. 4D), suggesting that increased cell-cycle- and proliferation-associated transcriptional signatures were not accompanied by a corresponding increase in the mitotic spindle signature.

**Figure 3.**
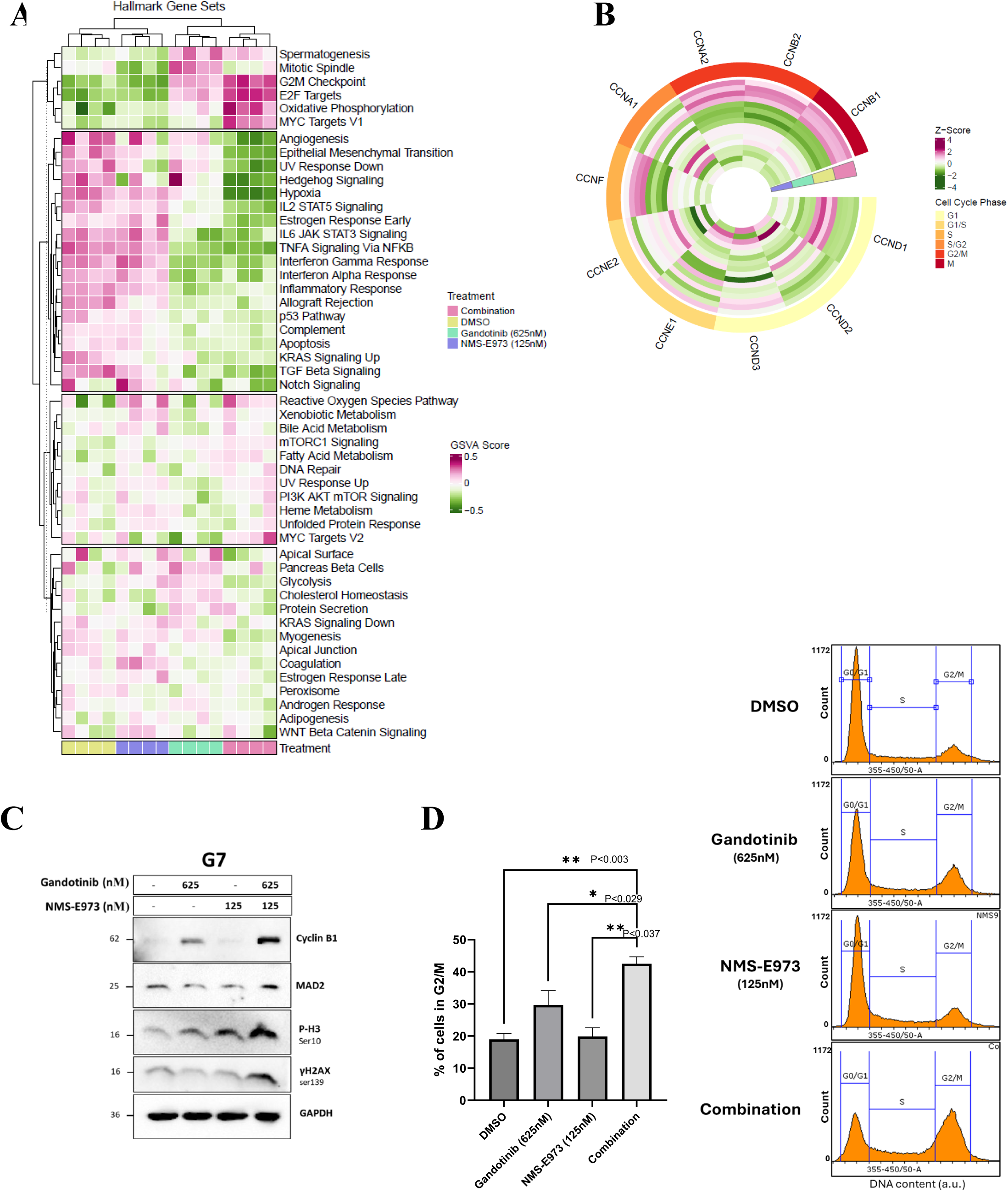
Combined JAK2 and HSP90 inhibition alters cell-cycle transcriptional programmes and promotes G2/M accumulation in G7 spheroids. **(A)** RNA-seq analysis of G7 spheroids following 24 h treatment with DMSO, Gandotinib (625nM), NMS-E973 (125nM) or the combination. Heatmap shows gene set variation analysis (GSVA) scores across 50 MSigDB Hallmark gene sets, with treatment groups indicated. **(B)** Circular heatmap showing z-scored expression of detected cyclin genes across treatment conditions. Cyclins are annotated according to their associated cell-cycle phase. **(C)** Western blot analysis of G7 spheroids following 24 h treatment with the indicated compounds. Cyclin B1, MAD2, phospho-Histone H3 Ser10 and γH2AX were assessed, with GAPDH shown as a loading control. **(D)** Flow-cytometric cell-cycle analysis of G7 spheroids following 24 h treatment under the same conditions as in (C). The percentage of cells in G2/M is shown as mean ± SD; n=2 technical and 3 biological replicates; adjusted *P* values are indicated.. Statistical significance in all the graphs was assessed using one-way ANOVA followed by Dunnett’s multiple comparisons test. Adjusted *P* values are indicated on the graph.

We next examined expression of individual cyclins to determine whether these transcriptional changes were associated with particular stages of the cell cycle. Combination treatment increased expression of cyclins associated with later stages of the cell cycle, including Cyclin A2, Cyclin B2 and Cyclin B1, whereas earlier cell-cycle cyclins such as Cyclin D1 were not similarly increased (Fig. 3B). Together, these transcriptional changes were consistent with accumulation of GBM stem cells in later stages of the cell cycle and altered progression through G2/M. We therefore examined proteins associated with mitotic progression in G7 spheroids. After 24 h combination treatment, Cyclin B1, a regulator of G2/M progression, MAD2, a spindle-assembly checkpoint protein, and phospho-Histone H3 Ser10, a marker of mitotic chromatin, were increased (Fig. 3C). We also observed increased γH2AX, a marker of DNA damage signalling that can also increase during mitosis.^24^ Flow-cytometric analysis also showed an increased proportion of cells in G2/M following combination treatment (Fig. 3D). Together, these transcriptomic, protein and DNA-content analyses support altered cell-cycle regulation, G2/M accumulation and changes in mitotic-associated signalling in G7 spheroids following combination treatment.

### Combined JAK2 and HSP90 inhibition alters mitotic regulation across heterogeneous patient-derived GBM cell lines

To investigate whether the mechanistic responses observed in G7 were also seen across different GBM molecular backgrounds, we examined three patient-derived GBM cell lines, GCGR-E13, GCGR-E31 and GCGR-E57, previously characterised as classical, proneural and mesenchymal subtypes, respectively.^5^ We used complementary approaches to examine mitotic morphology, γH2AX accumulation and cell-cycle progression in these GBM lines. In GCGR-E13 cells, immunofluorescence staining for α-tubulin and DAPI was used to examine mitotic spindle and chromosome organisation. Combination treatment increased the proportion of cells showing abnormal mitotic morphology, including altered chromosome and spindle organisation (Fig. 4A). This was accompanied by increased MAD2 and γH2AX, although Cyclin B1 did not show a combination-specific increase. In GCGR-E31 cells, combination treatment increased mean nuclear γH2AX fluorescence intensity, as assessed by quantitative immunofluorescence (Fig. 4B). Western blot also showed increased MAD2 and phospho-Histone H3 Ser10, while Cyclin B1 was reduced following NMS-E973 and combination treatment. These results showed that the pattern of individual mitotic markers differed between cell lines, but changes in mitotic-associated markers and increased γH2AX were common features of the combination response. To follow cell-cycle changes over time, we used GCGR-E57 cells stably expressing the fluorescent ubiquitination-based cell-cycle indicator (FUCCI), in which Cdt1 marks G1 and Geminin marks S/G2/M phases.^25^ Combination treatment of E57 FUCCI cells induced a progressive increase of TagGFP2-Geminin-positive cells, consistent with accumulation in S/G2/M-associated cell-cycle states (Fig. 4C). This was accompanied by increased Cyclin B1, phospho-Histone H3 Ser10 and γH2AX after 24 h treatment. A similar FUCCI response was observed when NMS-E973 was replaced with the alternative HSP90 inhibitor Pimitespib (Supplementary Fig. 5), suggesting that the phenotype was not restricted to a single HSP90 inhibitor.

**Figure 4.**
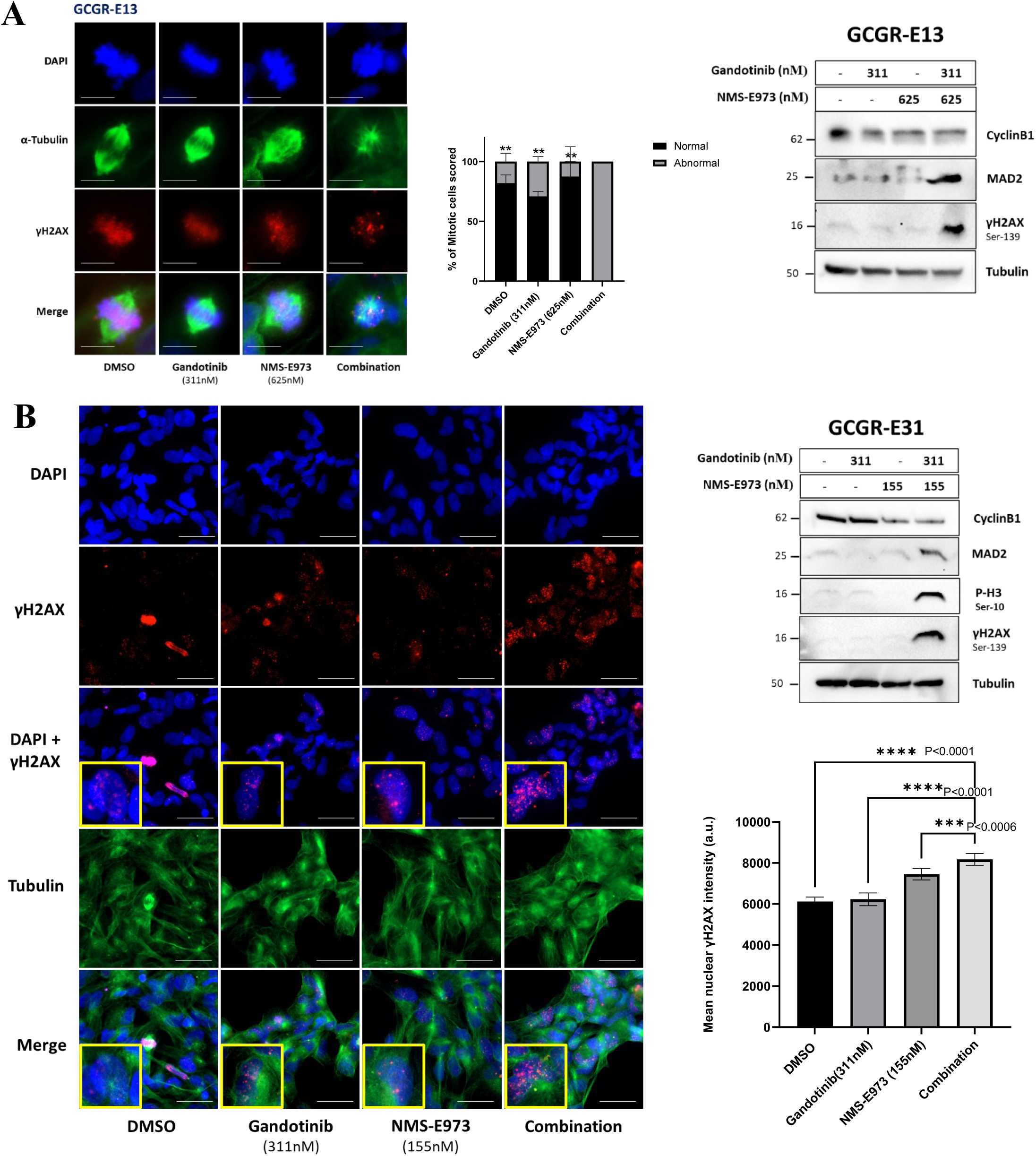
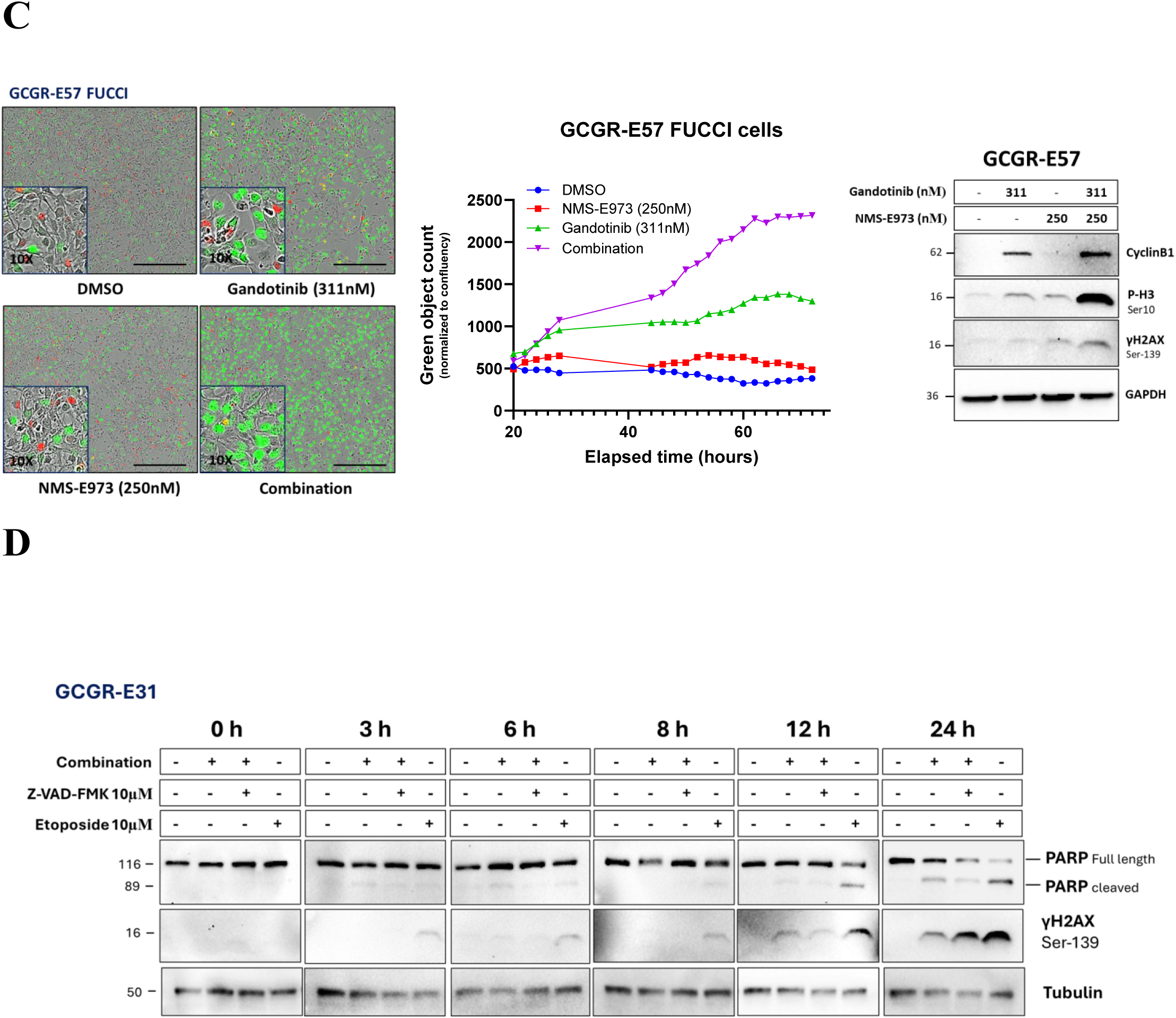
Combined JAK2 and HSP90 inhibition alters mitotic and cell-cycle regulation across heterogeneous patient-derived GBM models. Complementary mechanistic responses were examined in GCGR-E13 (classical), GCGR-E31 (proneural) and GCGR-E57 (mesenchymal) GBM cell lines. **(A)** Left: representative immunofluorescence images of GCGR-E13 cells following 24 h treatment with the indicated compounds. α-Tubulin, green; γH2AX, red; DAPI, blue. Scale bar, 10 μm. Middle: quantification of normal and abnormal mitotic morphology based on chromosome and spindle organisation, expressed as a percentage of total mitotic cells scored in each treatment group, n=2 technical and three biological replicates. Right: expression of Cyclin B1, MAD2 and γH2AX under the same treatment conditions. α-Tubulin is shown as a loading control. **(B)** Left: representative immunofluorescence images of GCGR-E31 cells following 24 h treatment with the indicated compounds. γH2AX, red; α-tubulin, green; DAPI, blue. Scale bar, 50 μm. Middle: mean nuclear γH2AX fluorescence intensity (a.u.), n=2 technical and three biological replicates. Right: expression of Cyclin B1, MAD2, phospho-Histone H3 Ser10 and γH2AX under the same treatment conditions. α-Tubulin is shown as a loading control. **(C)** Left: representative live-cell images of GCGR-E57 FUCCI cells following 72 h treatment with the indicated compounds. Middle: green FUCCI-positive object count normalised to cell confluency over time. Right: expression of Cyclin B1, phospho-Histone H3 Ser10 and γH2AX following 24 h treatment. GAPDH is shown as a loading control. **(D)** Time course expression of full-length and cleaved PARP and γH2AX in GCGR-E31 cells treated with DMSO, the combination (Gandotinib 311nM+NMS-E973 155nM), the combination plus z-VAD-fmk (10 μM), or etoposide (10 μM) for 0, 3, 6, 8, 12 and 24 h. α-Tubulin is shown as a loading control. Statistical significance in all the graphs was assessed using one-way ANOVA followed by Dunnett’s multiple comparisons test. Adjusted *P* values are indicated on the graph.

To further investigate the γH2AX response observed in GCGR-E31 cells, we performed a time-course experiment to determine whether γH2AX accumulation occurred early after treatment or developed later, using etoposide as a DNA-damaging control (Fig. 4D). Etoposide induced γH2AX at early timepoints, whereas γH2AX following combined JAK2 and HSP90 inhibition appeared later and was most evident at the later timepoints. Addition of the pan-caspase inhibitor z-VAD-fmk reduced PARP cleavage but did not prevent γH2AX accumulation, suggesting that the delayed γH2AX response was not simply caused by caspase-dependent apoptotic DNA fragmentation. Together, these complementary findings support altered cell-cycle and mitotic regulation following combined JAK2 and HSP90 inhibition across patient-derived GBM models representing different molecular backgrounds, accompanied by increased γH2AX. These observations extend the mechanistic responses initially identified in G7 spheroids.

### Combined JAK2 and HSP90 inhibition reduces GBM tumour burden in cerebral organoid co-cultures

We next tested the combination in a more complex 3D model using human cerebral organoid–GBM co-cultures (Supplementary Fig. 6A). GFP-Luciferase-labelled G7 cells were co-cultured with human iPSC-derived cerebral organoids and treated with DMSO, Gandotinib, NMS-E973 or combination for two weeks. Tumour burden was assessed using complementary imaging and histological approaches. Multiphoton and live-cell imaging showed extensive GFP-positive tumour areas in control organoids, whereas combination treated organoids showed a clear reduction in GFP positive area, which was confirmed by quantification (Fig. 5A).

**Figure 5.**
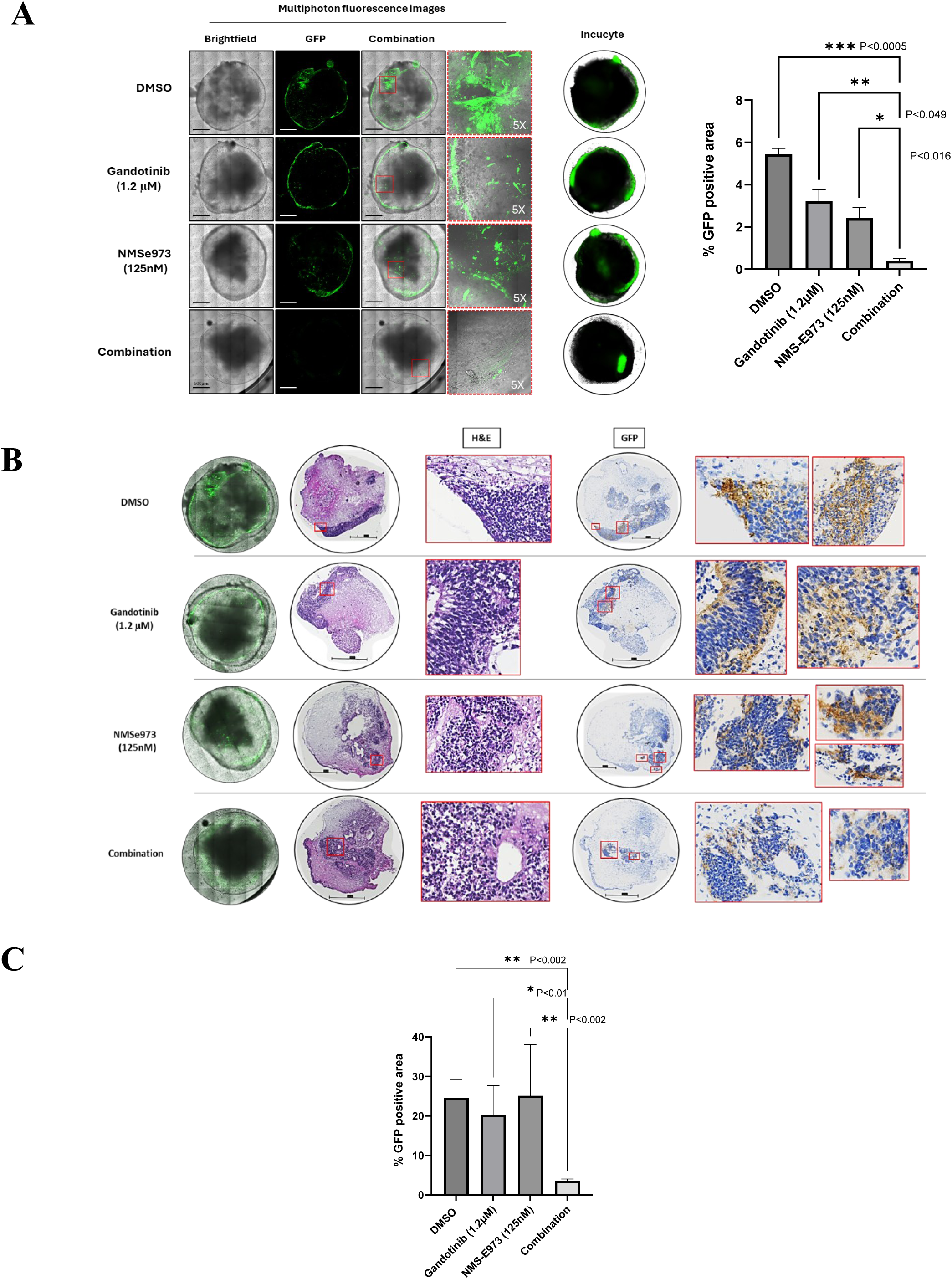
JAK2 and HSP90 co-inhibition reduces tumour burden in GBM–cerebral organoid co-cultures. **(A)** Left: representative multiphoton fluorescence and IncuCyte images of GFP-Luc-labelled G7 GBM–cerebral organoid co-cultures treated with DMSO, NMS-E973 (125 nM), Gandotinib (1.2 μM), or the combination. Right: quantification of GFP-positive area in the corresponding organoids (n = 2 technical and three biological replicates). Scale bar, 500 μm.**(B)** Representative H&E staining and GFP immunohistochemistry of control and drug-treated GBM–cerebral organoid sections. Scale bar, 500 μm.**(C)** Quantification of GFP-positive tumour area in organoid sections (n = 5 organoids in each technical replicate of three independent experiments). Statistical significance was assessed using one-way ANOVA followed by Dunnett’s multiple comparisons test. Adjusted *P* values are indicated on the graph.

We next examined the organoids by histology to determine whether this reduction was also evident within the organoids. H&E staining provided an overview of the organoid tissue architecture, while GFP immunohistochemistry identified G7 tumour cells and showed a reduction in GFP-positive tumour area following combination treatment (Fig. 5B). Quantification of GFP positive tumour area showed a significant reduction in the combination treated organoids compared with the other treatment groups (Fig. 5C). To provide another measure of the remaining tumour burden, organoid co-cultures were dissociated after two weeks of treatment and luciferase activity was measured from the GFP-Luc-labelled G7 cells (Supplementary Fig. 6B, C). Luciferase activity was significantly reduced following combination treatment compared with DMSO and the single agent conditions (Supplementary Fig. 6C). Since luciferase signal was derived specifically from the labelled G7 cells, this provided an additional measure of tumour cells remaining within the organoids after treatment and was consistent with the imaging and histological findings. Together, these three approaches showed a consistent reduction in GBM tumour burden within a 3D human GBM cerebral organoid co-culture model following combined JAK2 and HSP90 inhibition.

We also examined the effect of combined JAK2 and HSP90 inhibition in non-malignant human neural and glial cell types, including primary human foetal astrocytes, hESC-derived cortical neurons and iPSC-derived microglia, using GCGR-E13 cells as a patient-derived GBM comparator under the same drug-response assay conditions (Supplementary Fig. 7A-E). Cells were treated with NMS-E973 in combination with increasing concentrations of Gandotinib, and cell number was assessed by automated quantification of Hoechst-positive nuclei following 72 h treatment. Across the concentrations tested, cell number remained relatively stable in astrocytes, cortical neurons and microglia, whereas GCGR-E13 cells showed concentration dependent reduction (Supplementary Fig. 7B-E). Representative immunofluorescence images confirmed GFAP-positive astrocytes, MAP2-positive cortical neurons and Iba1-positive microglia-like cells maintain their cell differentiation markers and morphological phenotypes across the treatment conditions. These results show that the non-malignant human neural and glial cell types tested were less sensitive than GBM cells across the same drug concentration range.

### Combined JAK2 and HSP90 inhibition reduces tumour burden in orthotopic GBM xenografts

We next tested the combination in an orthotopic GBM model. GFP-Luc-labelled G7 cells were implanted intracranially into CD1 nude mice and, after tumour formation and IVIS based randomisation, mice were allocated to vehicle, Pacritinib (100 mg/kg), NMS-E973 (30 mg/kg) or combination treatment groups, with each compound administered according to its respective dosing schedule, as detailed in the Methods. Mice were culled 1 week after the final treatment (Fig. 6A). Complete details of the dosing regimen are described in the Methods. Histological analysis confirmed intracranial tumour growth in all treatment groups by H&E staining and immunohistochemistry using a human mitochondria antibody. Tumours from combination-treated mice appeared smaller and less extensive than those in vehicle or single agent treated mice (Fig. 6A). Quantification of tumour area showed a significant reduction in tumour burden following combination treatment compared with vehicle and either single agent (Fig. 6B).

**Figure 6.**
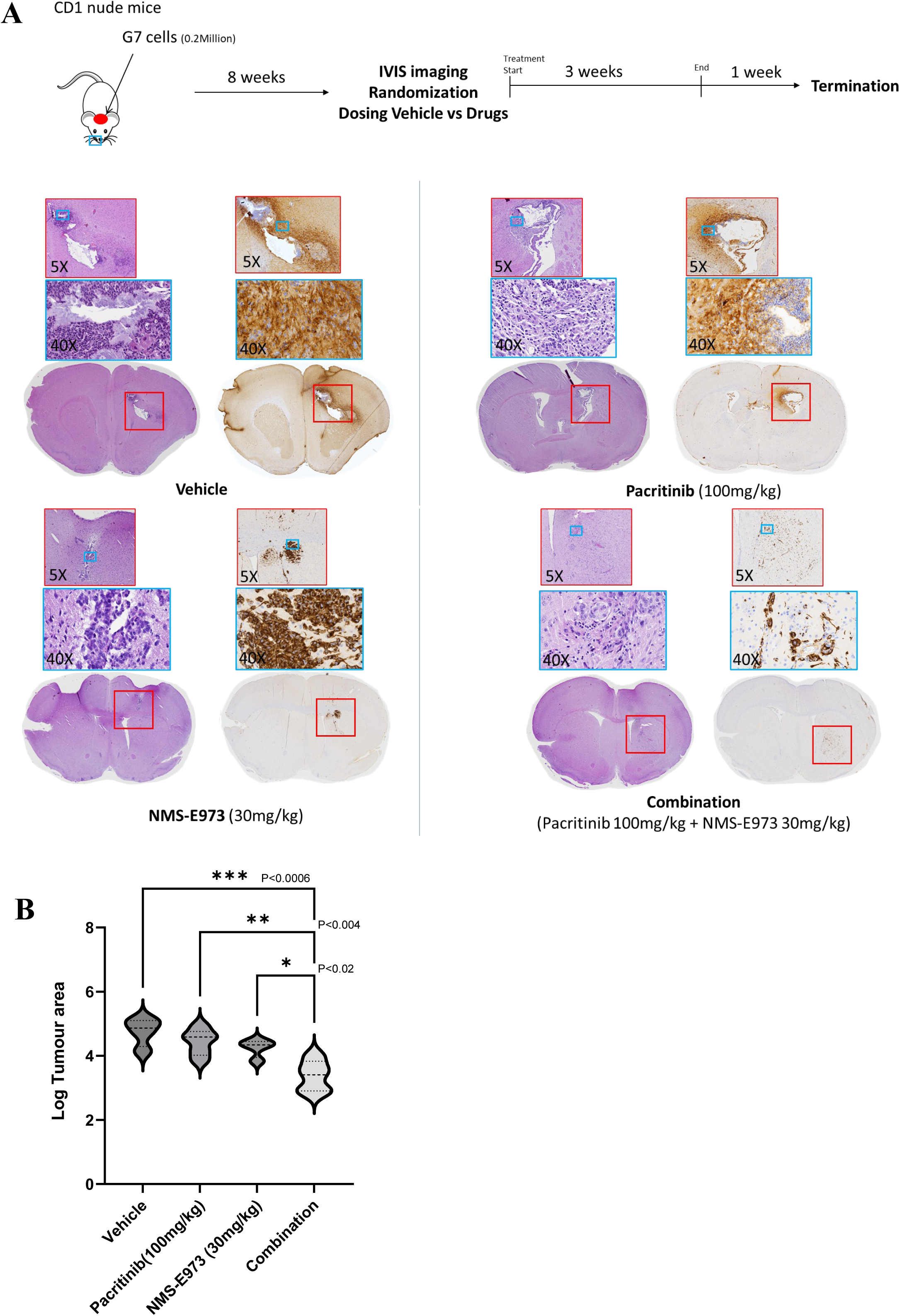
Combined JAK2 and HSP90 inhibition reduces tumour burden in an orthotopic G7 xenograft model. **(A)** Top: schematic of the in vivo efficacy study. G7 GBM cells were implanted intracranially into CD1 nude mice. Following tumour establishment, mice were randomised into four treatment groups (n = 5 per group): vehicle, Pacritinib (100 mg/kg), NMS-E973 (30 mg/kg), or the combination. Pacritinib was administered orally once daily, five days per week for three weeks (15 doses), while NMS-E973 was administered intraperitoneally using a three-days-on/one-day-off schedule for a total of 12 doses over 15 days. Mice were culled 1 week after the final treatment. Details of the dosing regimen are provided in methods. Bottom: representative H&E and human mitochondria immunohistochemistry from each treatment group. Low-magnification whole-brain sections and corresponding higher-magnification regions are shown. **(B)** Quantification of tumour area from histological sections (n = 5 mice per group), shown as log tumour area. Statistical significance was assessed using one-way ANOVA followed by Dunnett’s multiple comparisons test. Adjusted *P* values are indicated on the graph.

Because intracranial tumour growth can vary across different regions of the brain, we also examined additional sections from individual mice to assess the distribution and extent of tumour growth more broadly. Representative H&E and human mitochondria-labelled sections from individual mice are shown in Supplementary Fig. 8A. Across these additional sections, vehicle and single agent treated mice generally showed more extensive tumour growth, whereas tumour burden was reduced in the combination treated group.

Body weight was monitored throughout treatment as a general measure of treatment tolerability. Most of the mice-maintained body weight close to their starting weight during the dosing period. One mouse in the Pacritinib treated group showed a fall in relative body weight to below 0.9, but this occurred only towards the end of treatment and was not representative of the group as a whole. No consistent reduction in body weight was observed in the combination treated mice (Supplementary Fig. 8B). To provide supporting pharmacodynamic information, we also carried out a separate short PD study using the orthotopic 005-GSC mouse model. These murine GBM stem cells carry activated H-Ras and AKT signalling on a p53+/− background and can be grown as syngeneic intracranial tumours in C57BL/6 mice.^26^ Pacritinib treatment was associated with reduced p-STAT3 Y705 staining, while NMS-E973 treatment altered p-ERK and p-S6 Ser235/236 staining within tumour bearing regions (Supplementary Fig. 9). These findings provide supporting evidence of drug associated pathway modulation within intracranial tumour tissue in a separate GBM model. Together, these results show that combined Pacritinib and NMS-E973 treatment reduces tumour burden in an orthotopic G7 xenograft model.

## Discussion

In this study, we identify combined JAK2 and HSP90 inhibition as a broadly effective pharmacological strategy across heterogeneous glioblastoma (GBM) models. Importantly, synergistic interactions were observed in 12 of the 13 GBM cell lines tested, with maximum ZIP scores above 10 for at least one JAK2/HSP90 inhibitor pair (Fig. 1C). Only GCGR-E34 remained within the additive range. This widespread response is particularly encouraging given the molecular and phenotypic heterogeneity of GBM, which presents a major challenge for therapies targeting individual signalling pathways.^5^ Previous studies have demonstrated anti-tumour activity following inhibition of either JAK2/STAT3 or HSP90 in GBM, ^27–30^ while combined targeting of these pathways has shown synergistic effects in JAK2-dependent haematological malignancies.^15–17^ Our findings extend these observations by demonstrating that this pharmacological interaction is reproducible across different patient-derived GBM backgrounds and multiple inhibitor pairs. Furthermore, combination activity was maintained in increasingly complex models, including 3D spheroids, cerebral organoid co-cultures and an orthotopic GBM xenograft model, supporting the potential relevance of this strategy beyond conventional in vitro cell culture.

The interaction between JAK2 and HSP90 inhibition may reflect the interconnected nature of their signalling networks. JAK2 is an established HSP90 client protein, and HSP90 inhibition has previously been shown to destabilise JAK2 and suppress downstream JAK–STAT signalling in other cancer models.^15^ In our study, NMS-E973 induced a rapid increase in STAT3 Y705 phosphorylation, which was reduced by the addition of Gandotinib. Similar adaptive JAK–STAT signalling responses have been reported following HSP90 inhibition in other cancer settings.^20,21,31^ Although these findings suggest possible compensatory signalling, we have not established the upstream mechanism responsible for STAT3 phosphorylation or its contribution to drug synergy. The broad activity of HSP90 inhibitors against multiple signalling pathways suggests that the enhanced combination response may involve additional survival mechanisms beyond STAT3 regulation alone. Combined JAK2 and HSP90 inhibition also enhanced apoptotic responses across several GBM cell lines, as demonstrated by caspase-3 activation and PARP cleavage. These findings are consistent with previous reports of apoptosis following JAK2/STAT3 or HSP90 inhibition ^27,32,33^ and suggest that enhanced apoptotic cell death contributes to the broad anti-tumour activity of the combination.

Our transcriptomic, protein and flow-cytometric analyses suggest that combined JAK2 and HSP90 inhibition disrupts normal G2/M regulation and mitotic progression. Complementary experiments in GCGR-E13, GCGR-E31 and GCGR-E57 cells, representing classical, proneural and mesenchymal GBM backgrounds,^5^ respectively, further supported this response, including abnormal mitotic morphology and accumulation in later cell-cycle states. Previous studies have shown that HSP90 inhibition can interfere with key regulators of G2/M progression, while STAT3-associated signalling has also been implicated in cell-cycle regulation in GBM.^32,34–36^ Together, these findings support a shared disturbance of cell-cycle and mitotic regulation across different GBM models, although they do not establish a uniform mitotic arrest.

An additional finding was increased γH2AX following combination treatment in G7 and the three GCGR cell lines examined. Although γH2AX is commonly used as a marker of DNA double-strand breaks, H2AX phosphorylation can also occur during mitosis without externally induced DNA damage.^24,37^ Furthermore, prolonged or abnormal mitosis has been associated with secondary DNA-damage signalling and genomic stress.^38,39^ In our E31 time-course experiments, etoposide induced an early γH2AX response, whereas γH2AX accumulation following combined JAK2 and HSP90 inhibition developed later. Inhibition of caspase activity using z-VAD-fmk reduced PARP cleavage but did not prevent γH2AX accumulation, indicating that the response was not simply explained by caspase-dependent apoptotic DNA fragmentation. Together with the observed changes in mitotic-associated markers and cell-cycle distribution, these findings suggest an association between altered mitotic progression and γH2AX accumulation. However, they do not distinguish mitosis-associated H2AX phosphorylation from DNA damage or broader chromatin stress arising during abnormal cell-cycle progression.

The anti-tumour activity of the combination was further supported by experiments using human cerebral organoid–GBM co-cultures. These models provide a more complex, brain-like cellular environment in which to examine tumour growth and treatment response than conventional monolayer cultures.^40,41^ In our study, combined treatment reduced growth of G7 cells within cerebral organoids, with consistent findings from fluorescence imaging, GFP immunohistochemistry and luciferase-based measurements. The agreement between these complementary approaches strengthens the evidence for reduced growth of tumour cells within the organoid environment. Importantly, non-malignant human astrocytes, cortical neurons and microglia showed relatively limited changes in cell number or morphology at drug concentrations that markedly reduced GCGR-E13 GBM cell numbers. This differential sensitivity provides encouraging preliminary evidence of a potentially favourable response profile, although differences in cellular proliferation and the short-term nature of these assays mean that a therapeutic window cannot yet be fully established.

Importantly, the anti-tumour response was also observed *in vivo*. Combined Pacritinib and NMS-E973 treatment significantly reduced histologically assessed tumour burden in orthotopic G7 xenografts compared with vehicle or either single agent. Body weight remained generally stable during combination treatment, with no consistent treatment-associated reduction observed in this group. The choice of these compounds for intracranial evaluation was supported by previous studies reporting brain exposure and anti-tumour activity of Pacritinib in GBM models and NMS-E973 in an intracranial tumour model.^9,11^ We also obtained supporting pharmacodynamic evidence in a separate syngeneic 005-GSC intracranial model, where short-course Pacritinib treatment was associated with reduced phospho-STAT3 Y705 staining and NMS-E973 altered phospho-ERK and phospho-S6 staining within tumour-bearing regions. These findings are consistent with modulation of relevant signalling pathways within intracranial tumour tissue. Together, the *in vivo* findings extend the activity observed in cell-based and organoid models and support further evaluation of this combination strategy in GBM.

The present study has three main limitations. First, the *in vivo* efficacy experiment was designed to assess tumour burden rather than survival, and it remains to be determined whether the reduction in intracranial tumour growth translates into prolonged survival. Second, our findings are based predominantly on pharmacological inhibition, and complementary genetic approaches would help establish the specific contributions of JAK2 and HSP90 to the observed interaction. Finally, although increased γH2AX was a reproducible feature of the combination response and occurred alongside altered cell-cycle and mitotic regulation, its precise biological basis remains unresolved. Further studies will be needed to distinguish mitosis-associated H2AX phosphorylation from chromatin stress or DNA damage arising during abnormal mitotic progression. Overall, our findings identify combined JAK2 and HSP90 inhibition as a promising therapeutic strategy that produces widespread synergistic activity across heterogeneous GBM models. The consistent anti-tumour responses observed in 2D and 3D cultures, cerebral organoid co-cultures and an orthotopic xenograft model, together with evidence of altered cell-cycle regulation, mitotic progression and apoptotic cell death, provide a strong rationale for further investigation of this combination as a potential treatment option for GBM patients.

## Material and methods

### Cell culture

Patient-derived glioblastoma stem cell lines were obtained from the Glioma Cellular Genetics Resource (GCGR), Edinburgh. The GCGR models used in this study included GCGR-E13, E15, E20, E21, E28, E31, E34, E55, E57, E63, L3 and L5. Cells were maintained in DMEM/Ham’s F12 medium (Merck Life Sciences, #D8437) supplemented with glucose (8 mM; Merck Life Sciences, #G8644), β-mercaptoethanol (100 μM; Thermo Fisher Scientific, #31350-010), MEM non-essential amino acids (Thermo Fisher Scientific, #11140-035), BSA (0.015%; Thermo Fisher Scientific, #15260-037), recombinant basic FGF (12 ng/mL; PeproTech, #100-18B), recombinant EGF (12 ng/mL; PeproTech, #315-09), B27 (1:100; Thermo Fisher Scientific, #17504-044), N2 (1:200; Thermo Fisher Scientific, #17502-048) and laminin (4–10 μg/mL; R&D Systems, #3446-005-01), as previously described.^5,42^ The patient-derived G7 glioblastoma stem cell model was also used throughout the study and was maintained under serum-free stem cell culture conditions as previously described.^43^ Cells were cultured at 37 °C in a humidified atmosphere containing 5% CO₂ and routinely passaged before reaching full confluence.

### Compound treatment and drug combination screening

The JAK2 inhibitors Gandotinib, Pacritinib and Fedratinib and the HSP90 inhibitors NMS-E973, Pimitespib, AUY922 and Ganetespib were used for single-agent and combination studies. Compounds were prepared in DMSO. For single-agent dose–response experiments, cells were plated in 384-well plates and allowed to attach or establish spheroids before compound treatment. Compounds were tested over a concentration range spanning their respective dose–response profiles, and cell survival was assessed by counting Hoechst-stained nuclei after 72 h treatment. IC₅₀ values were calculated from concentration–response curves and used to guide the concentration ranges selected for subsequent combination experiments.

Drug combinations were tested using 7 × 7 concentration matrices containing the corresponding single-agent treatment arms, following the approach used previously^42^. Compounds were dispensed using a D300e digital dispenser (Tecan), with the final DMSO concentration kept below 0.5% and constant across treatment and control wells. Combination matrices were centred around concentrations producing measurable single-agent responses to allow assessment of drug interactions across a range of effect levels. Cell viability was normalised to the corresponding DMSO treated controls. Experiments were performed in at least three independent biological replicates.

### Cell viability and dose–response analysis

For two-dimensional drug-response assays, GBM cells were seeded into 384-well plates at densities appropriate for each cell line and allowed to establish before treatment. Cells were exposed to serial dilutions of JAK2 or HSP90 inhibitors for 72 h. At the end of treatment, cell survival was quantified by nuclei counting following Hoechst staining. Cell numbers were normalised to the corresponding DMSO treated controls and expressed as percentage survival. Dose response curves were generated and IC₅₀ values calculated by nonlinear regression using a four-parameter logistic model (log[inhibitor] versus response, variable slope) in GraphPad Prism version 10 (GraphPad Software, Boston, MA, USA). IC₅₀ values were subsequently converted to pIC₅₀values for comparison of compound potency across the GBM cell-line panel.

For G7 spheroid dose response experiments, spheroids were generated under non-adherent conditions and treated with increasing concentrations of indicated compounds for 72 h. Spheroid viability was assessed using PrestoBlue reagent (Thermo Fisher Scientific, Paisley, Scotland, UK, #A13262) according to the manufacturer’s instructions, and fluorescence values were normalised to DMSO-treated controls. IC₅₀ values were calculated from the resulting concentration response curves and converted to pIC₅₀ (−log₁₀ IC₅₀ in molar units) for each independent experiment.

### Human astrocyte, microglia and cortical neuron drug-response assays

Primary human fetal astrocytes (Caltag Medsystems, SC-1800) were seeded at 2,500 cells per well in poly-D-lysine-coated 384-well plates (Greiner Bio-One, 781946) in Human Astrocyte Media (Caltag Medsystems, SC1801). After 2 days, a half-medium change was performed and cells were treated with DMSO control (0.1% final concentration) or JAK2/HSP90 inhibitor combinations across the indicated dose ranges using a TECAN Fluent automated liquid handler. Cells were maintained for a further 3 days before fixation and staining. Human iPSC-derived myeloid precursors (Kolf 2C) were generated as previously described^44^ and seeded at 5,000 cells per well in gelatin- and poly-D-lysine-coated 384-well plates. Cells were differentiated for 12 days in Advanced DMEM/F12 supplemented with 1% N2, 2 mM GlutaMAX, 50 μM β-mercaptoethanol, 1% penicillin/streptomycin, 100 ng/mL IL-34 and 10 ng/mL GM-CSF, with half-medium changes every 2–3 days. Following the medium change on day 12, cells were treated with DMSO or the indicated drug combinations for 3 days before fixation.

Cortical neurons were generated from the RC17 human ESC line (RCe021-A) using a previously described dual-SMAD inhibition protocol with modifications.^45^ Neural precursor cells were differentiated towards cortical neurons and maintained in neuronal differentiation medium as described in the original protocol. For drug response experiments, vehicle or indicated drug combinations were added following final medium change, and cells were cultured for a further 3 days before fixation.

For all three cell types, cells were fixed in 4% paraformaldehyde and processed for immunocytochemistry. Astrocytes were identified using GFAP (Agilent, Z0334; 1:1000), microglia using Iba1 (Abcam, ab5076; 1:2000), and cortical neurons using MAP2 (Abcam; 1:1000). Nuclei were stained with Hoechst 33258 (1:5000). Images were acquired using an ImageXpress Micro Confocal system (Molecular Devices), with nine fields captured per well using a 20× objective. Image analysis was performed using MetaXpress software. Hoechst-positive nuclei were segmented using the Find Round Objects module, and the mean nuclei count across the nine fields was used as the viability readout for each well.

### Drug synergy analysis

Drug interactions were analysed using SynergyFinder (version 3.0)^46^. Normalised cell survival data from 7 × 7 dose–response matrices, including the corresponding single-agent concentrations, were converted to percentage inhibition and analysed using the zero-interaction potency (ZIP) model. ZIP synergy scores were calculated across each concentration matrix, with positive scores indicating a greater response than expected under the ZIP reference model. A ZIP score >10 was considered indicative of synergy. For comparison of JAK2 and HSP90 inhibitor combinations across GBM models, the maximum ZIP synergy score from each dose matrix was reported together with the percentage inhibition observed at the corresponding concentration pair. Synergy landscapes and dose–response matrices were generated using SynergyFinder^46^. Combination experiments were performed across a minimum of three biological replicates.

### Three-dimensional spheroid invasion assay

GBM cells were seeded at 2000 cells per well in 96-well ultra-low-attachment U-bottom plates (Corning Life Sciences, #7007) in normal growth medium and centrifuged at 200 × g. Cells were allowed to form a single spheroid per well over 3–4 days before being overlaid with Matrigel (Thermo Fisher Scientific, #11543620) at a final concentration of 4 mg/mL. Spheroids were subsequently treated with the indicated concentrations of JAK2 and HSP90 inhibitors, either alone or in combination. Spheroid growth and invasive outgrowth were monitored every 3 h for up to 12 days using an IncuCyte S3 live-cell imaging system (Sartorius) with phase and brightfield imaging using a 10× objective. Images were analysed using the IncuCyte 3D Spheroid Invasion analysis module. Invasive area was quantified as the brightfield object area extending beyond the spheroid core. Three independent experiments were performed, with two spheroids per treatment condition in each experiment.

### Live-cell analysis of caspase-3 activation

Caspase-3 activation following drug treatment was monitored by live-cell imaging using BioTrac 530 Red Caspase-3 Dye. G7 spheroids were treated with DMSO, Gandotinib, NMS-E973 or the combination in the presence of the caspase-3 reporter and imaged over time using an IncuCyte live-cell imaging system. Caspase-3 activation was quantified as integrated red fluorescence intensity from each spheroid. The assay was also performed across a panel of patient-derived GBM cell lines (E13, E28, E21, E57 and E31). Cells were treated with DMSO, the indicated concentrations of Gandotinib or NMS-E973, or their combination and imaged every 3 h for 40–80 h, depending on the cell line. Caspase-3 activity was quantified as integrated red fluorescence intensity per image and plotted over time. Three independent experimental replicates were used for each treatment condition.

### Western blotting

Cells or spheroids were collected following treatment at the indicated timepoints, washed with PBS and lysed in ice-cold RIPA buffer (50 mM Tris-HCl, pH 8.0, 150 mM NaCl, 1% Triton X-100, 0.5% sodium deoxycholate and 0.1% SDS) supplemented with phosphatase and protease inhibitor cocktails (PhosSTOP and complete ULTRA; Roche). Samples were incubated on ice for 30 min with gentle agitation and centrifuged at high speed to remove insoluble material. Protein concentration in the resulting supernatant was determined using Coomassie Plus Protein Assay Reagent (Thermo Fisher Scientific), and samples were normalised for protein content. Equal amounts of protein were resolved on 4–15% Mini-PROTEAN TGX gels (Bio-Rad) and transferred onto PVDF membranes. Membranes were blocked before overnight incubation with primary antibodies at 4 °C. Following washing with PBS-T, membranes were incubated with the appropriate HRP-conjugated secondary antibodies for 1 h at room temperature, washed again and visualised by enhanced chemiluminescence using a ChemiDoc imaging system (Bio-Rad). Western blotting was used to assess pharmacodynamic pathway modulation following JAK2 and HSP90 inhibition, including phospho-STAT3 Y705, STAT3, phospho-AKT S473, AKT, phospho-ERK1/2 and ERK1/2. Apoptotic responses were assessed using full-length and cleaved PARP, while Cyclin B1, MAD2, phospho-Histone H3 Ser10 and γH2AX were examined to investigate treatment-associated cell-cycle and mitotic responses. GAPDH or α-tubulin were used as loading controls as indicated. Details of primary antibodies and working dilutions are provided in Supplementary Table 1.

### RNA sequencing and pathway analysis

G7 spheroids were treated for 24 h with DMSO, Gandotinib, NMS-E973 or the combination before collection for RNA sequencing. Total RNA was isolated from four independent biological replicates and subjected to quality assessment prior to library preparation and sequencing. Total RNA samples were quantified using a Qubit 2.0 Fluorometer (Thermo Fisher Scientific, #Q32866) with the Qubit RNA Broad Range kit (#Q10210), assessed for DNA contamination using the Qubit dsDNA HS assay (#Q32854), and evaluated for integrity on an Agilent 2100 Bioanalyzer (Agilent Technologies, #G2939AA) using RNA 6000 Nano chips (#5067-1511). Libraries were prepared from 500 ng RNA per sample using the Lexogen QuantSeq 3′ mRNA-Seq V2 Library Prep Kit (FWD) for Illumina with 12-nt unique dual indexes (Lexogen GmbH, A1-191.96), following the manufacturer’s protocol. Following first- and second-strand cDNA synthesis, libraries were amplified for 16 cycles and purified using magnetic beads. Library concentration and fragment size were assessed using the Qubit dsDNA HS assay and Agilent Bioanalyzer DNA HS kit (#5067-4626), respectively, and libraries were pooled equimolarly at 1 nM. Sequencing was performed on an Illumina NextSeq 2000 platform (#20038897) using a NextSeq 1000/2000 P1 XLEAP-SBS Reagent Kit, 100 cycles (#20100983), with one P1 flow cell in a single-end 1 × 100-cycle format; base-call data were converted to FASTQ files using Illumina BaseSpace. Read counts were normalised and filtered using DESeq2 (v1.48.2)^47^, gene set variation analysis was performed using GSVA (v2.2.1)^22^, and heatmaps were plotted using ComplexHeatmap (v2.24.1)^48^ using the R (v4.5.1) software for statistical computing.

### Flow cytometric analysis of cell-cycle distribution

G7 spheroids were treated for 24 h with DMSO, Gandotinib, NMS-E973 or the combination. Following treatment, spheroids were collected, dissociated with accutase (Sigma-Aldrich; #A6964) to obtain a single-cell suspension and washed twice with ice-cold PBS. Cells were resuspended in PBS and fixed by addition to ice-cold 70% ethanol while gently vortexing, followed by incubation at 4 °C for at least 2 h. On the day of analysis, fixed cells were washed with PBS and allowed to rehydrate for 15 min at room temperature. Equal numbers of cells (1–2 × 10⁶ per sample) were resuspended in 300 µL DAPI staining solution containing 1 µg/mL DAPI and 0.1% Triton X-100 in PBS and incubated for 30 min at room temperature protected from light. DAPI fluorescence was acquired on a BD LSRFortessa™ flow cytometer using 405 nm excitation and a 450/50 emission filter in linear mode. DNA-content profiles were analysed using Flowing Software version 2.5.1. An untreated asynchronous sample was used to define the positions of the G1 and G2 peaks, and the proportions of cells in G0/G1, S and G2/M phases were determined from the resulting DNA-content distributions.

### Immunofluorescence staining and imaging

For immunofluorescence analysis of GBM cells, GCGR-E13 and GCGR-E31 cells were seeded onto coverslips coated with 1 μg/mL laminin-I (R&D Systems; #3446-005-01) and allowed to adhere before treatment with DMSO, Gandotinib, NMS-E973 or the combination for 24 h. Following treatment, cells were fixed by direct addition of formaldehyde to a final concentration of 4% for 15 min at 37 °C. Cells were washed with Tris-buffered saline (TBS) and incubated with 0.1 M glycine in TBS for 10 min at room temperature to quench residual formaldehyde. Cells were then permeabilised with 0.1% Triton X-100 in TBS for 5 min and blocked in 2% BSA containing 0.1% Triton X-100 in TBS for 1 h at room temperature. Primary antibodies against α-tubulin and γH2AX were diluted in blocking buffer and incubated overnight at 4 °C. Cells were washed three times and incubated with the appropriate fluorescently labelled secondary antibodies for 45 min at room temperature in the dark. Following further washing, nuclei were counterstained with DAPI and coverslips were mounted using antifade mounting medium. Representative fluorescence images shown in Figure 4 were acquired using a Photometrics Prime BSI CMOS camera (Photometrics, Tucson, AZ, USA) fitted to a Zeiss AxioImager M2 fluorescence microscope with Plan-Apochromat objectives and a Zeiss Colibri 7 LED light source (Carl Zeiss, Cambridge, UK). Images were acquired using a 100× objective in Zeiss ZEN 3.5 software, with identical acquisition settings used across treatment conditions within each experiment. α-Tubulin staining was used to assess microtubule and mitotic spindle organisation. For GCGR-E13 cells, mitotic cells were classified as showing normal or abnormal mitotic morphology based on chromosome and spindle organisation, and the proportion of abnormal mitoses was calculated relative to the total number of mitotic cells analysed. For quantitative analysis of γH2AX in GCGR-E31 cells, images were acquired separately using an ImageXpress Micro Confocal system (Molecular Devices) and analysed using MetaXpress software. Nuclei were segmented using DAPI staining, and mean γH2AX fluorescence intensity was measured within the nuclear region. Measurements from individual fields were averaged within each independent biological experiment before statistical analysis.

For immunostaining of the non-malignant neural cell models, primary human astrocytes, iPSC-derived microglia and hESC-derived cortical neurons were fixed after 3 days of drug treatment using 4% paraformaldehyde in PBS. Cells were washed three times with PBS using an AquaMax 4000 automated plate washer (Molecular Devices) and stored in PBS at 4 °C until staining. Cells were permeabilised with 0.2% Triton X-100 in PBS for 10 min and blocked for 30 min using 6% normal goat serum for rabbit primary antibodies or 6% normal donkey serum for goat primary antibodies. Primary antibodies were diluted in the appropriate blocking buffer and incubated for 12 h at 4 °C. Astrocytes were stained using rabbit anti-GFAP (Agilent; #Z0334; 1:1000), microglia using goat anti-Iba1 (Abcam; #ab5076; 1:2000), and cortical neurons using Alexa Fluor 647-conjugated anti-MAP2 (Abcam; 1:1000).

Following primary antibody incubation, cells were washed three times with PBS. Astrocytes and microglia were then incubated for 1 h at room temperature with Alexa Fluor 488 goat anti-rabbit (#A32731TR, Thermo Fisher Scientific; 1:1000) or Alexa Fluor 647 donkey anti-goat (#A-11055, Thermo Fisher Scientific; 1:1000), respectively. Hoechst 33258 (Sigma-Aldrich; 1:5000) was included for nuclear staining. MAP2-stained cortical neuron cultures were incubated with Hoechst 33258 without additional secondary antibody staining. Cells were subsequently washed three times with PBS and stored in PBS at 4 °C before imaging. Images of astrocytes, microglia, cortical neurons and the corresponding GCGR-E13 comparison cultures were acquired using an ImageXpress Micro Confocal system (Molecular Devices) with a 20× objective. Nine fields were acquired per well and analysed using MetaXpress software. Hoechst-positive nuclei were segmented using the Find Round Objects module, and the mean nuclei count across the nine fields was used as the cell-number readout for each well.

### FUCCI live-cell imaging

GCGR-E57 cells stably expressing the fluorescent ubiquitination-based cell cycle indicator (FUCCI, in which Cdt1 marks G1 and Geminin marks S/G2/M phases)^25^ were seeded into 96-well plates and allowed to establish before drug treatment. Cells were treated with DMSO, Gandotinib, NMS-E973 or the combination and monitored by live-cell imaging using an IncuCyte Zoom microscope (Essen BioScience) maintained at 37 °C. Images were acquired at regular intervals over several days and analysed using the integrated IncuCyte analysis software. Cell-cycle progression was assessed by quantifying green FUCCI-positive objects over time. Green object counts were normalised to cell confluency to account for differences in cell number between treatment conditions. The same approach was used in experiments in which NMS-E973 was replaced with the alternative HSP90 inhibitor Pimitespib.

### Cerebral organoid–GBM co-culture and drug treatment

Human iPSC-derived cerebral organoids were differentiated using a modified Lancaster protocol ^49^ which included addition of 10µM SB431542 (Tocris) and 100nM LDN-193189 (APExBIO) for neural induction and addition of 20ng/ml BDNF and NT3 (Peprotech) to the medium at D30 for two weeks to aid neuronal maturation as previously reported.^49^ For tumour co-culture experiments, day-30 cerebral organoids were co-cultured with GFP-luciferase-labelled G7 GBM cells for approximately 2 weeks to allow establishment of the tumour–organoid cultures. Co-cultures were then treated for a further 2 weeks with DMSO, Gandotinib (1.2 μM), NMS-E973 (125 nM), or the combination. Tumour growth and distribution within the organoids were monitored during treatment using IncuCyte live-cell imaging and multiphoton fluorescence microscopy. GFP-positive tumour area was quantified from multiphoton images as a measure of tumour burden. At the experimental endpoint, organoid co-cultures were processed using complementary readouts. For histological analysis, organoids were fixed, paraffin embedded, sectioned and subjected to H&E staining and GFP immunohistochemistry. GFP-positive tumour area within the sections was quantified as a histological measure of tumour burden. In parallel, organoids allocated for biochemical assessment were enzymatically dissociated and lysed. Firefly luciferase activity from the GFP-Luc-labelled G7 cells was measured using a glow-based luciferase assay according to the manufacturer’s instructions. Luciferase values were normalised to the DMSO-treated control and used as an additional biochemical measure of tumour cell burden.

### Multiphoton microscopy

Two-photon fluorescence images were acquired using a custom-built multi-modal microscope setup.^50^ Briefly, a picoEmerald S (APE, Berlin, Germany) laser provided both a tunable pump laser (700-980 nm, 2 ps, 80 MHz repetition rate) and a spatially and temporally overlapped Stokes laser (1031 nm, 2 ps, 80 MHz repetition rate). The output beams were inserted into the scanning unit of an Olympus FV1000-MPE microscope using a series of dielectric mirrors and a 3× lens-based beam-expanding module. The resulting 3.6 mm beams were expanded by a further 3.6× within the microscope and directed into an Olympus XLPL25XWMP N.A. 1.05 objective lens using a short-pass 690 nm dichroic mirror (Olympus). Backscattered emission signals from two-photon fluorescence were separated from any backscattered excitation light using a short-pass 690 nm dichroic mirror and IR cut filter (Olympus). A series of filters and dichroic mirrors were then used to deconvolve the different emission signals onto one of 4 available photo-multiplier tubes (PMT). GFP two-photon fluorescence signals were filtered using the following series of filters: FF552-Di02, FF483/639-Di01 and FF510/84 (Semrock). Transmitted light images were collected by condenser lens and PMT. The pump laser was tuned to 797.2 nm, powers after the objective were measured at 30 mw for the pump and Stokes laser. All images were recorded at 512 × 512 pixels with a dwell time of 4 µs per pixel by FV10-ASW software (Olympus) using the MATL function for image tiling. Image analysis and processing was performed in ImageJ (Fiji) 1.54P.^51^

### Human astrocyte, microglia and cortical neuron drug-response assays

Primary human fetal astrocytes (Caltag Medsystems, SC-1800) were seeded at 2,500 cells per well in poly-D-lysine-coated 384-well plates (Greiner Bio-One, 781946) in Human Astrocyte Media (Caltag Medsystems, SC1801). After 2 days, a half-medium change was performed and cells were treated with DMSO control (0.1% final concentration) or JAK2/HSP90 inhibitor combinations across the indicated dose ranges using a TECAN Fluent automated liquid handler. Cells were maintained for a further 3 days before fixation and staining. Human iPSC-derived myeloid precursors (Kolf 2C) were generated as previously described^44^ and seeded at 5,000 cells per well in gelatin- and poly-D-lysine-coated 384-well plates. Cells were differentiated for 12 days in Advanced DMEM/F12 supplemented with 1% N2, 2 mM GlutaMAX, 50 μM β-mercaptoethanol, 1% penicillin/streptomycin, 100 ng/mL IL-34 and 10 ng/mL GM-CSF, with half-medium changes every 2–3 days. Following the medium change on day 12, cells were treated with DMSO or the indicated drug combinations for 3 days before fixation. Cortical neurons were generated from the RC17 human ESC line (RCe021-A) using a previously described dual-SMAD inhibition protocol with modifications.^45^ Neural precursor cells were differentiated towards cortical neurons and maintained in neuronal differentiation medium as described in the original protocol. For drug response experiments, vehicle or indicated drug combinations were added following final medium change, and cells were cultured for a further 3 days before fixation.

For all three cell types, cells were fixed in 4% paraformaldehyde and processed for immunocytochemistry. Astrocytes were identified using GFAP (Agilent, Z0334; 1:1000), microglia using Iba1 (Abcam, ab5076; 1:2000), and cortical neurons using MAP2 (Abcam; 1:1000). Nuclei were stained with Hoechst 33258 (1:5000). Images were acquired using an ImageXpress Micro Confocal system (Molecular Devices), with nine fields captured per well using a 20× objective. Image analysis was performed using MetaXpress software. Hoechst-positive nuclei were segmented using the Find Round Objects module, and the mean nuclei count across the nine fields was used as the viability readout for each well.

### Animal ethics and welfare

Animal experiments were conducted in accordance with UK Home Office regulations and the Animals (Scientific Procedures) Act 1986. The G7 and 005 orthotopic studies were performed under UK Home Office project licence PP7510272 with ethical approval from the University of Edinburgh Animal Welfare and Ethical Review Body (PL05-21). Mice were maintained at the Western General Hospital animal facility, University of Edinburgh, under standard institutional husbandry conditions. They were monitored twice daily for the first 48–72 h following surgery and daily thereafter, with body weight and general clinical condition monitored throughout the experiment. Particular attention was given to neurological signs, including ataxia, impaired balance and reduced responsiveness, together with changes in grooming, posture, mobility and body weight. Humane endpoints were applied in accordance with the approved project licence, and animals showing signs of neurological impairment or deteriorating clinical condition were humanely euthanised. The experimental protocol had a prospective moderate severity classification.

### Orthotopic G7 xenograft efficacy study

Female CD1 nude mice (8–12 weeks old; n = 20) were used to establish orthotopic G7 glioblastoma xenografts. Mice were anaesthetised using isoflurane and positioned in a stereotactic frame. Following a midline scalp incision, a small burr hole was made 0.6 mm anterior and 1.5 mm right lateral to bregma. A suspension of GFP-Luc-labelled G7 glioblastoma stem cells (2 × 10⁵ cells in 2 µL) was implanted intracranially at a depth of 2.5 mm using a Hamilton syringe connected to a syringe pump, as previously described.^42^ Following injection, the needle was left in position for 4 min before withdrawal, the burr hole was sealed with bone wax and the wound was closed using absorbable sutures. Buprenorphine was administered subcutaneously for perioperative analgesia, followed by carprofen in drinking water for at least 48 h. Tumours were allowed to establish over 8 weeks. Tumour growth was monitored using an IVIS Lumina S5 imaging system following subcutaneous injection of D-luciferin (150 mg/kg) under isoflurane anaesthesia. Following tumour establishment and IVIS-based randomisation, mice were allocated to four treatment groups (n = 5 per group): vehicle, Pacritinib (100 mg/kg), NMS-E973 (30 mg/kg), or the combination. Pacritinib was administered by oral gavage once daily, five days per week for three weeks (15 doses), following a previously reported regimen in GBM mouse models.^52^ NMS-E973 was administered intraperitoneally once daily using an intermittent schedule of three consecutive treatment days followed by one day without treatment, for a total of 12 doses over 15 days. This intermittent schedule was adapted from previous preclinical studies of NMS-E973,^11^ with modifications to the dose, administration route and treatment duration. At the experimental endpoint, mice were humanely euthanised by cervical dislocation. Brains were collected, fixed in formalin and processed for paraffin embedding. Intracranial tumour burden was assessed by histological analysis and quantification of human mitochondria-positive tumour area, as described below. Five mice per treatment group were included in the final histological analysis (n = 5 per group). Treatment allocation and subsequent experimental assessments, including histological tumour area quantification, were performed blinded to treatment group identity.

### *In vivo* pharmacodynamic analysis in the 005 orthotopic GBM model

GFP-labelled 005 GBM cells were implanted intracranially into C57BL/6 mice. Fifteen days after implantation, tumour-bearing mice were randomised to receive vehicle, Pacritinib (100 mg/kg) via P.O, or NMS-E973 (30 mg/kg) via I.P. Mice were treated for 3 consecutive days and brains were collected on day 18 for pharmacodynamic assessment. Brains were fixed, paraffin embedded and sectioned for H&E staining and immunohistochemistry. GFP staining was used to identify tumour-bearing regions. Sections from Pacritinib-treated mice were stained for phospho-STAT3 Y705 and phospho-ERK1/2, while sections from NMS-E973-treated mice were stained for phospho-ERK1/2 and phospho-S6 Ser235/236. Staining patterns within GFP-positive tumour regions were compared with corresponding vehicle-treated sections to assess pathway modulation following short-course drug treatment. 005-GSC cells are murine glioblastoma stem cells originally derived from gliomas generated by activated H-Ras and AKT signalling on a p53+/− background. Cells were maintained under serum-free neural stem-cell conditions as previously described.^26^ 005-GSCs form neurospheres *in vitro* and generate syngeneic intracranial tumours in C57BL/6 mice.

### Histology, immunohistochemistry and tumour-area quantification

At the experimental endpoint, mouse brains were collected, fixed in formalin overnight and processed for paraffin embedding. Coronal sections encompassing the tumour-bearing region were cut at 4 µm thickness and mounted onto glass slides. Sections were either stained with haematoxylin and eosin (H&E) or processed for immunohistochemistry (IHC).

For IHC, sections were dewaxed in xylene and subjected to antigen retrieval using 10 mM citrate buffer in a pressure cooker. Following washing, endogenous peroxidase activity was blocked using Dako REAL Peroxidase-Blocking Solution, followed by serum-free protein blocking. Sections were incubated overnight with primary antibody at 4 °C. Human G7 tumour cells within mouse brain tissue were identified using an anti-human mitochondria antibody (1:100; Bio-Techne, #NBP2-32980). The following day, sections were incubated with DAKO EnVision HRP secondary reagent and staining was visualised using DAB. Sections were counterstained with haematoxylin, dehydrated and mounted using DPX. Whole-brain sections were scanned at 40× magnification using a Hamamatsu NanoZoomer digital slide scanner S360 (Hamamatsu Photonics). Tumour area was quantified from the resulting whole-slide images using ImageJ (Fiji) 1.54P. Human mitochondria-positive tumour regions were manually identified and measured, and tumour area was used to compare tumour burden between treatment groups.

For cerebral organoid experiments, fixed organoids were paraffin embedded and sectioned before H&E staining and GFP immunohistochemistry. GFP-positive regions were used to identify G7 tumour cells within organoid sections, and total GFP-positive tumour area was quantified using ImageJ as an independent histological measure of tumour burden.

### Statistical analysis

Statistical analyses were performed using GraphPad Prism v10.2.0 (GraphPad Software, San Diego, CA, USA). Data are presented as mean ± SD unless otherwise stated. The number of independent biological replicates used for each experiment is indicated in the corresponding figure legend. For comparisons involving multiple treatment groups, one-way ANOVA followed by an appropriate multiple-comparisons test was used. Where specified, Dunnett’s multiple-comparisons test was applied to compare individual treatment groups with the combination-treated group. For the caspase-3 live-cell assay, statistical analysis was performed at the 48 h timepoint rather than across the full time course. Tumour-area measurements from the *in vivo* study were log-transformed before statistical analysis where indicated. Adjusted P values are shown in the corresponding figures. A P-value < 0.05 was considered statistically significant.

## Author Contributions

M.F., R.J.R.E., and N.O.C. conceived and co-ordinated the project, designed and interpreted the experiments. M.F., R.J.R.E., A.E.P.L, W.K.C, R.M, M.L., M.M, F.L, J.C.D., R.C., B.T.S., P.B., P.W.K.N., A.F.M., C.D., G.M.M., and V.S.M., performed the experiments. N.O.C., M.C.F., V.G.B., S.M.P., and J.O.M., contributed to supervision. M.F., R.J.R.E., A.E.P.L, W.K.C, R.M. and N.O.C. analysed the data. M.F., R.J.R.E., A.E.P.L., and N.O.C. prepared figures. M.F., R.J.R.E., A.E.P.L, W.K.C, R.M, M.L., M.M, F.L, J.C.D., R.C., B.T.S., P.B., contributed to methodology. M.F., and N.O.C., wrote the manuscript. All authors read and approved the manuscript.

