## Supplementary figures for "Combined JAK2 and HSP90 inhibition produces synergistic anti-tumour activity across preclinical glioblastoma models"

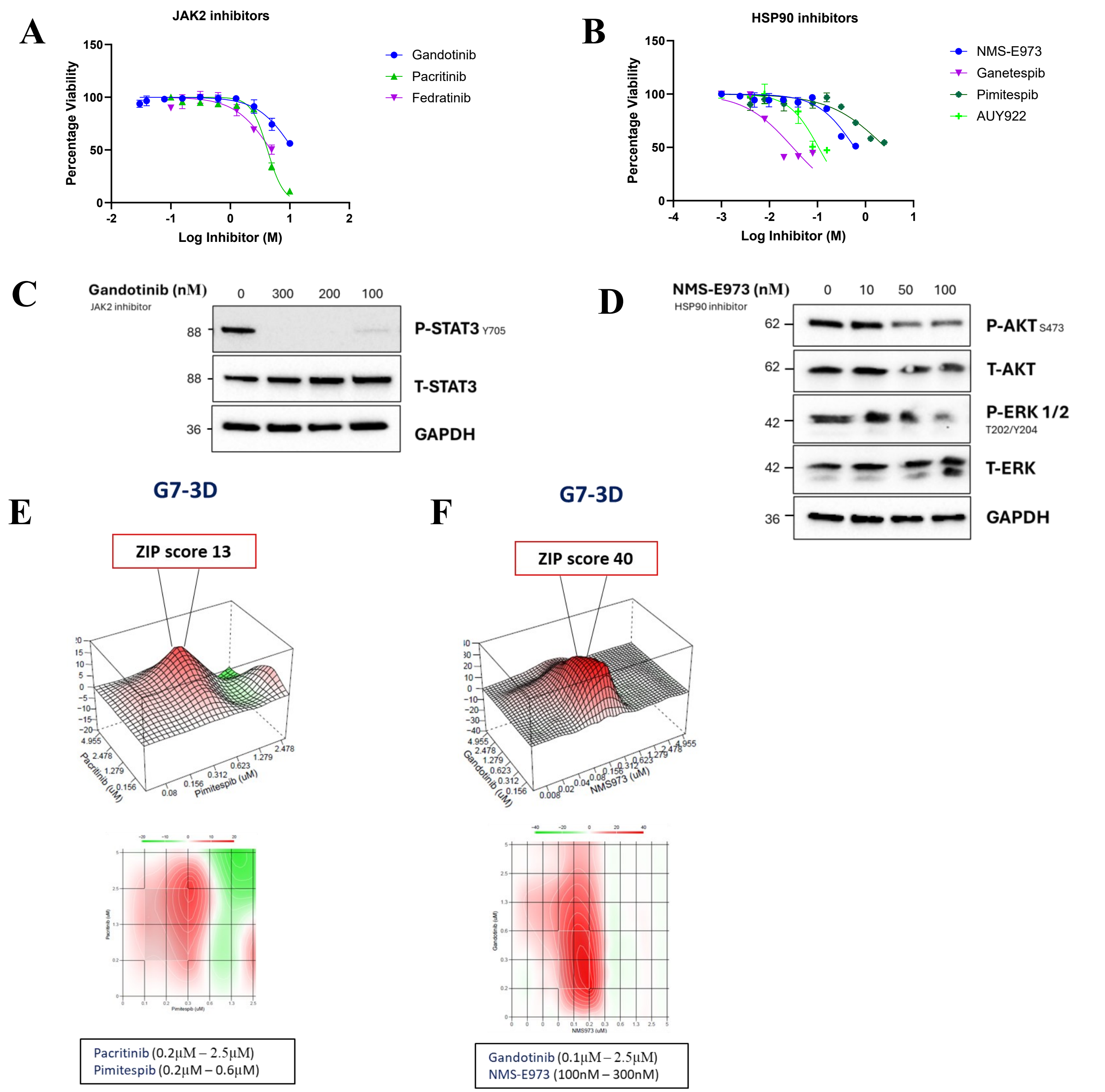


Pacritinib + Pimitespib

| Dose pair | Pacritinib (µM) | Pimitespib (µM) | ZIP synergy score | Loewe synergy score | Bliss synergy score | HSA synergy score |
|---|---|---|---|---|---|---|
| 1 | 2.5 | 0.312 | 13.01 | -5.52 | 16.96 | 14.94 |
| 2 | 1.25 | 0.312 | 11.60 | 0.64 | 11.90 | 9.86 |
| 3 | 2.5 | 0.155 | 9.92 | -5.44 | 8.87 | 5.95 |
| 4 | 2.5 | 0.08 | 7.47 | -3.87 | 3.75 | 0.50 |

Gandotinib + NMS-E973

| Dose pair | Gandotinib (µM) | NMS-E973 (µM) | ZIP synergy score | Loewe synergy score | Bliss synergy score | HSA synergy score |
|---|---|---|---|---|---|---|
| 1 | 0.311 | 0.125 | 24.53 | 20.20 | 30.41 | 25.27 |
| 2 | 0.623 | 0.125 | 38.72 | 31.50 | 39.03 | 34.22 |
| 3 | 1.25 | 0.125 | 54.79 | 1.10 | 58.04 | 53.18 |
| 4 | 1.25 | 0.063 | 21.38 | 14.97 | 10.69 | 11.85 |

**Supplementary Figure 1. Dose–response and target-engagement validation of JAK2 and HSP90 inhibitors in G7 spheroids. (A–B)** Dose–response curves for **(A)** JAK2 inhibitors Gandotinib, Pacritinib and Fedratinib, and **(B)** HSP90 inhibitors NMS-E973, Pimitespib, AUY922 and Ganetespib in G7 3D spheroids. Cell viability was measured using PrestoBlue following 72 h treatment. $IC_{50}$ values were determined from the dose–response curves and converted to $pIC_{50}$ for presentation in Figure 1B. Data are shown as mean ± SD from three independent experiments (n = 3). **(C-D)** Western blot analysis validating concentration-dependent modulation of downstream signalling pathways in G7 cells treated for 24 hours with **(C)** Gandotinib (monitoring downstream phospho-STAT3Y705 and total STAT3) and **(D)** NMS-E973 (monitoring client pathway readouts phospho-AKTS473, total AKT, phospho-ERK-P44/42-T202/Y204, and total ERK). GAPDH served as the loading control. **(E-F)** ZIP synergy landscapes, dose–response matrices and synergy scores at different concentrations for **(E)** Pacritinib + Pimitespib and **(F)** Gandotinib + NMS-E973 in G7 3D spheroids. **(G)** Western blot validation of pathway modulation at synergistic Gandotinib and NMS-E973 concentration pairs in G7 cells. Right panel: quantification of normalized P-STAT3 Y705 expression relative to DMSO, mean ± SD, n=2 technical and 3 biological replicates.**(H)** Time course western blot of p-STAT3(Y705) and total STAT3 in G7 cells treated with indicated drugs for indicated timepoints. GAPDH served as loading control. Right panels: quantification of normalized P-STAT3 Y705 expression relative to DMSO after 2 and 3 hours of treatment, mean ± SD, n=2 technical and 3 biological replicates. Statistical significance was assessed using one-way ANOVA followed by Dunnett's multiple comparisons test. Adjusted *P* values are indicated on the graph.

G

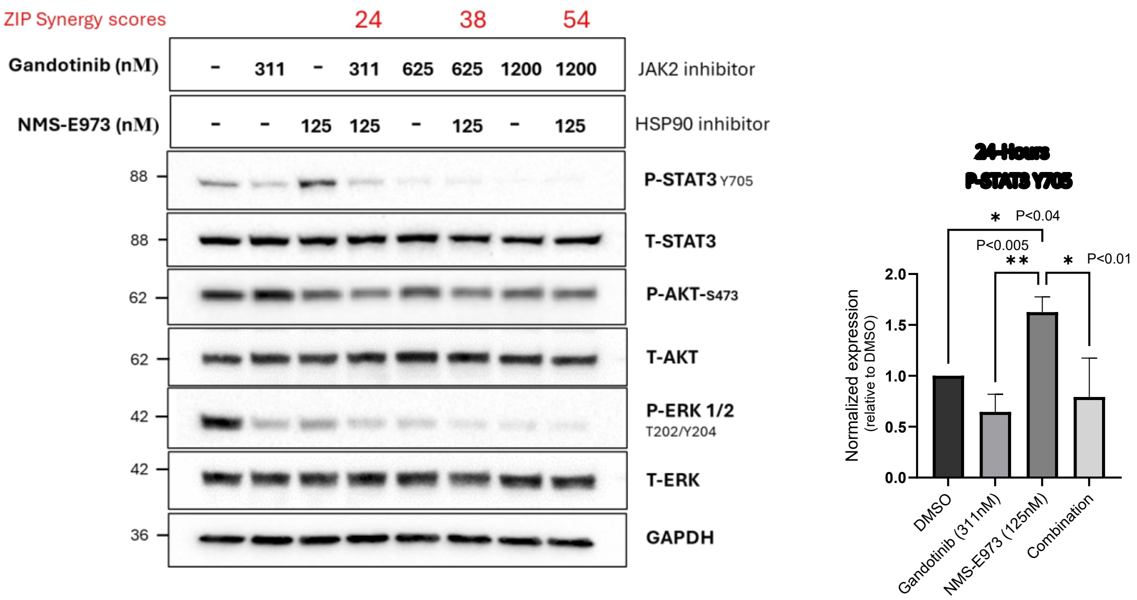
ZIP Synergy scores
24
38
54
Gandotinib (nM)
- 311 - 311 625 625 1200 1200
JAK2 inhibitor
NMS-E973 (nM)
- - 125 125 - 125 - 125
HSP90 inhibitor
88
P-STAT3 Y705
88
T-STAT3
62
P-AKT-S473
62
T-AKT
42
P-ERK 1/2
T202/Y204
42
T-ERK
36
GAPDH
24-Hours
P-STAT3 Y705
P<0.04
P<0.005
P<0.01
Normalized expression
(relative to DMSO)
DMSO
Gandotinib (311nM)
NMS-E973 (125nM)
Combination

H

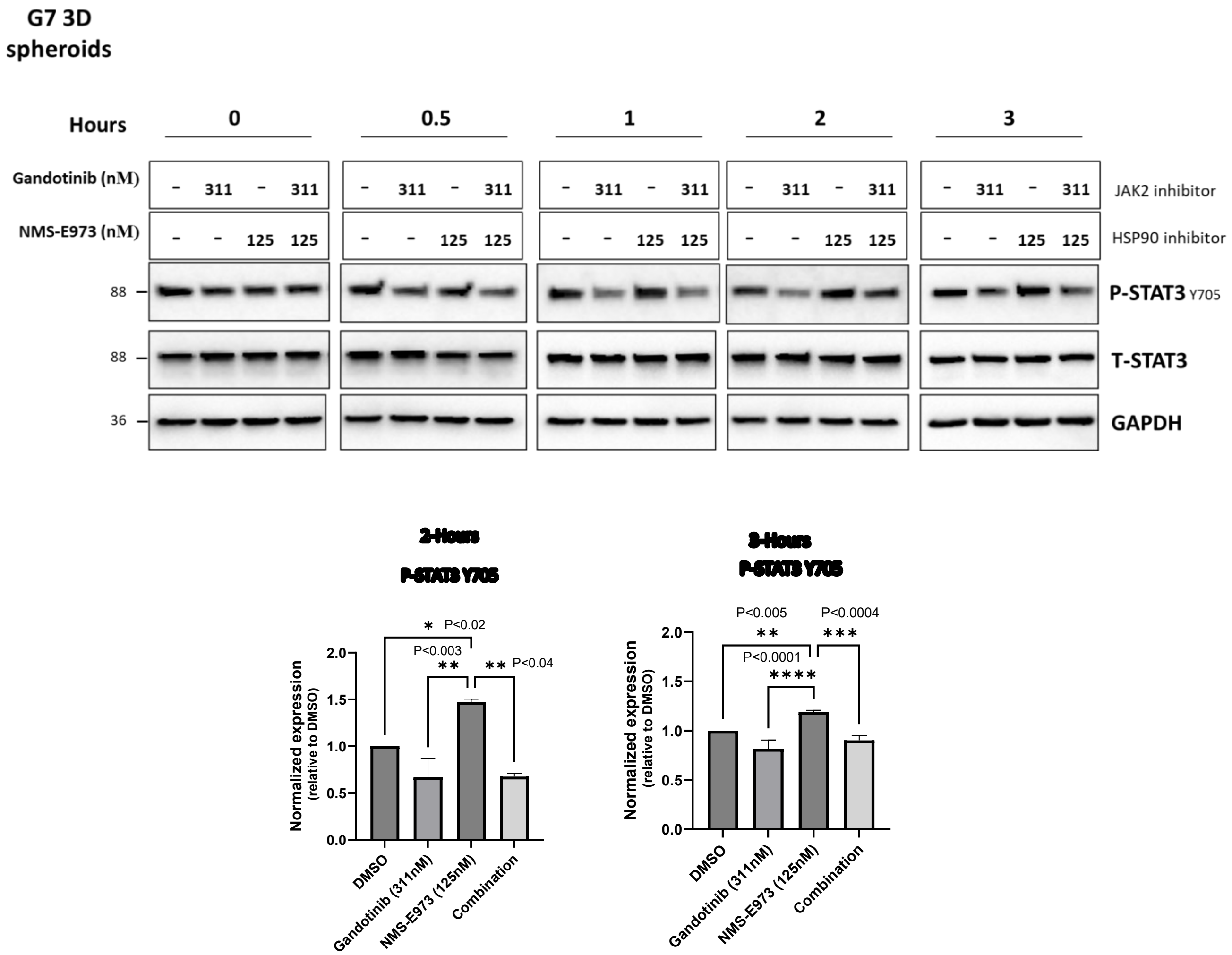
G7 3D
spheroids
Hours
0
0.5
1
2
3
Gandotinib (nM)
- 311 - 311
JAK2 inhibitor
NMS-E973 (nM)
- - 125 125
HSP90 inhibitor
88
P-STAT3 Y705
88
T-STAT3
36
GAPDH
2-Hours
P-STAT3 Y705
P<0.02
P<0.003
P<0.04
Normalized expression
(relative to DMSO)
DMSO
Gandotinib (311nM)
NMS-E973 (125nM)
Combination
3-Hours
P-STAT3 Y705
P<0.005
P<0.0004
P<0.0001

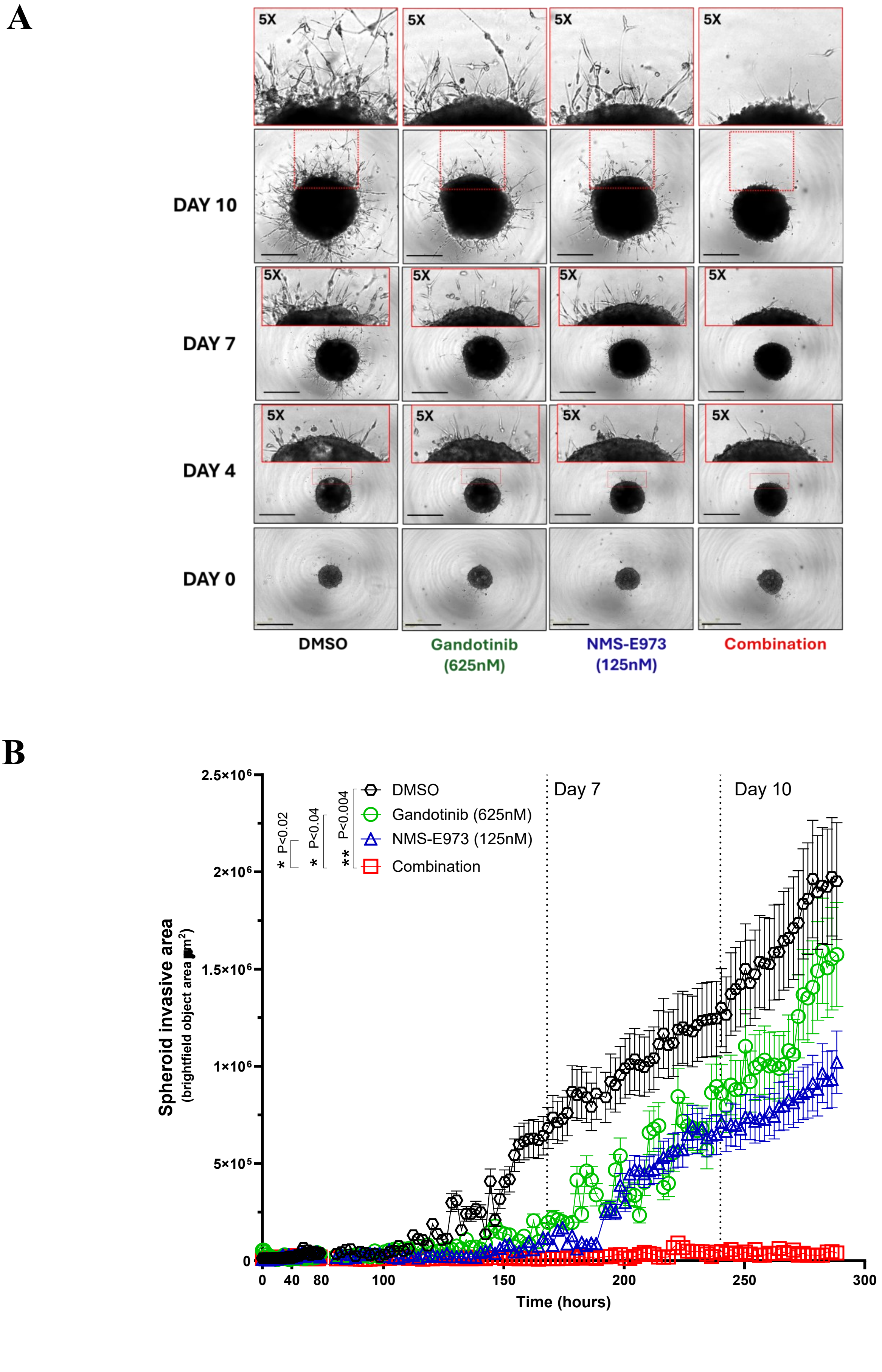


**Supplementary Figure 2: JAK2 and HSP90 co-inhibition suppresses invasive outgrowth of Matrigel-embedded G7 spheroids.** **(A)** Representative brightfield images of G7 GBM spheroids embedded in Matrigel and treated with the indicated compounds. Images were taken at Day 0,4,7 and10 using IncuCyte S3 live-cell imaging system (Sartorius). Enlarged regions (5x) show invasive outgrowth at spheroid–matrix interface. Scale bar represents 800 µm. **(B)** Quantification of invasive outgrowth area over 12 days. The measured area represents the brightfield object area extending beyond spheroid core. Statistical significance was assessed at day 10 (240h) timepoint using one-way ANOVA with Dunnett's multiple comparisons test; P values are indicated on the graph. Data are mean ± SD (n = 2 technical replicates per condition in each of three independent experiments).

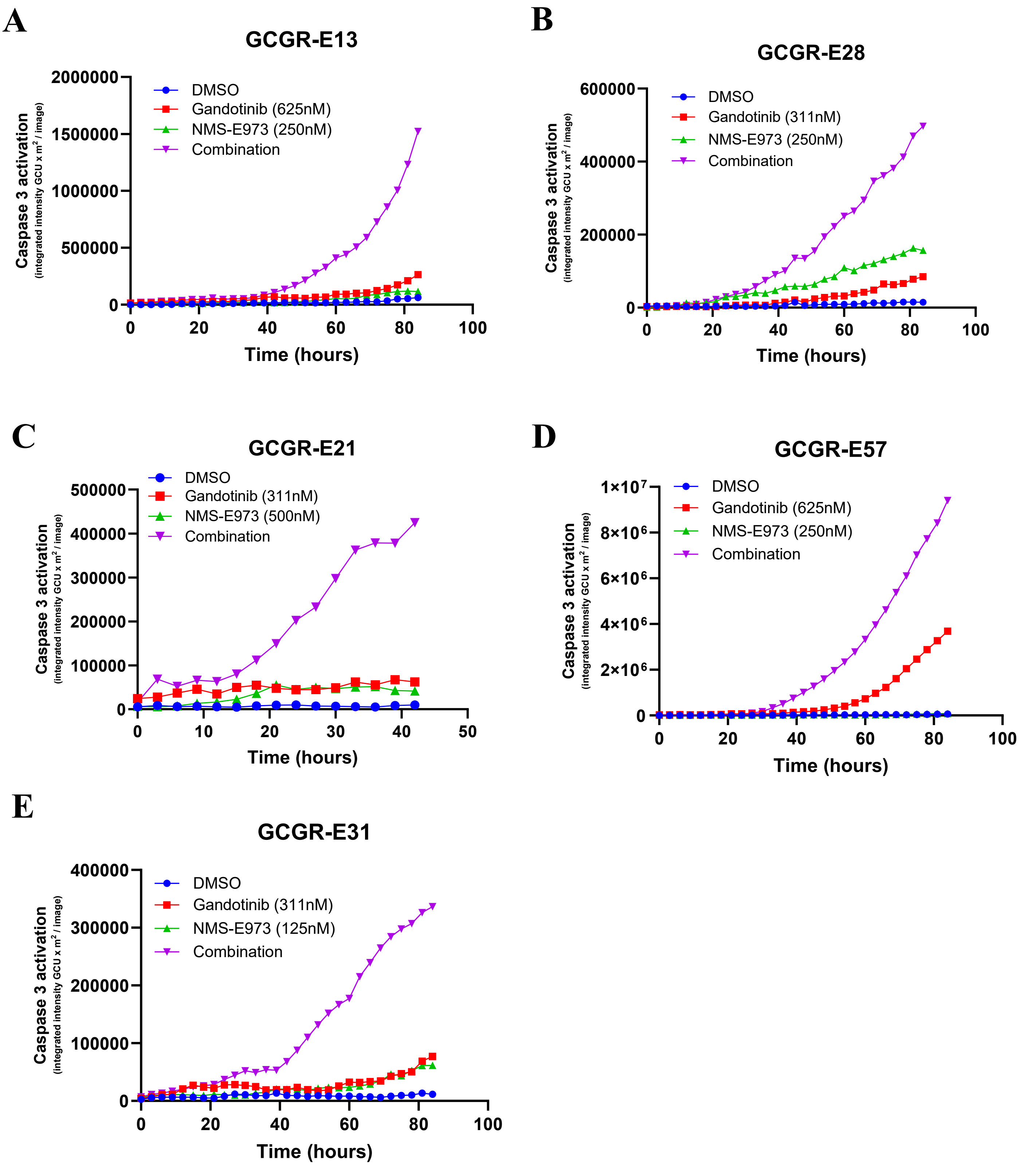


**Supplementary Figure 3. Live-cell quantification of caspase-3 activation across GBM lines treated with JAK2 and HSP90 inhibitors. (A–E)** Time-course quantification of caspase-3 activation in GBM cell lines treated with DMSO, Gandotinib, NMS-E973, or the combination. Caspase-3 activation was monitored by live-cell imaging every 3 h over 40–80 h and quantified as integrated fluorescence intensity per image. Panels show responses in GCGR- **(A)** E13, **(B)** E28, **(C)** E21, **(D)** E57 and **(E)** E31 cells.

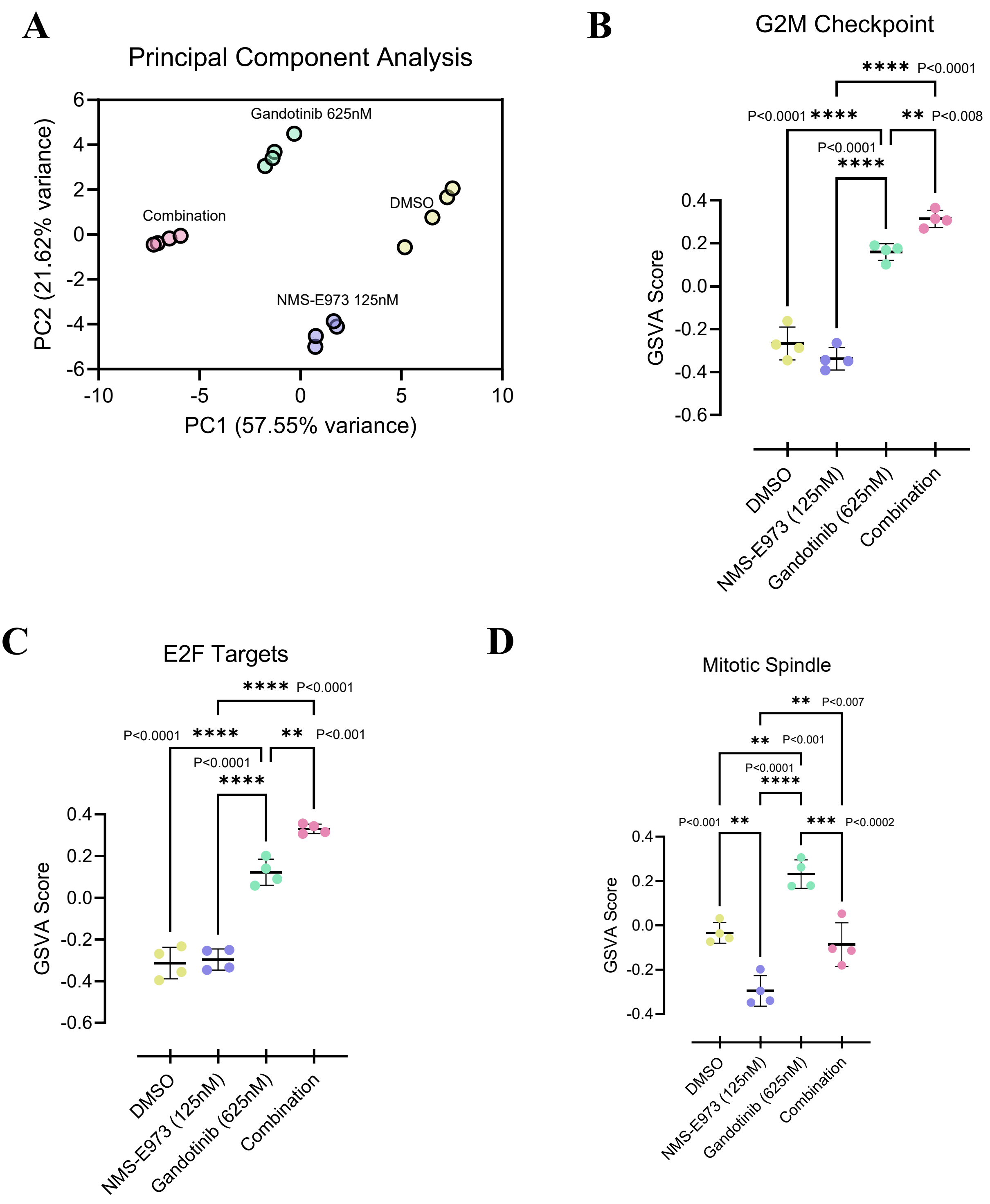


**Supplementary Figure 4.** RNA-seq analysis of cell-cycle associated transcriptional changes in G7 spheroids following JAK2 and HSP90 inhibition. **(A)** Principal component analysis of RNA-seq samples from G7 spheroids following 24 h treatment with DMSO, Gandotinib (625nM), NMS-E973 (125nM) or the combination. **(B-D)** GSVA scores for **(B)** G2/M checkpoint **(C)** E2F targets gene set and **(D)** Mitotic Spindle gene set across the indicated treatment groups. Individual replicates are plotted, (n = 4 per treatment group) with mean ± SD indicated; statistical significance was assessed by one way ANOVA followed by Tukey's multiple comparisons test are indicated on the graphs. Adjusted *P* values are indicated on the graphs.

# A

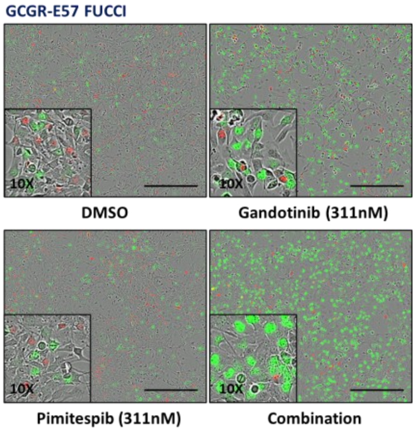


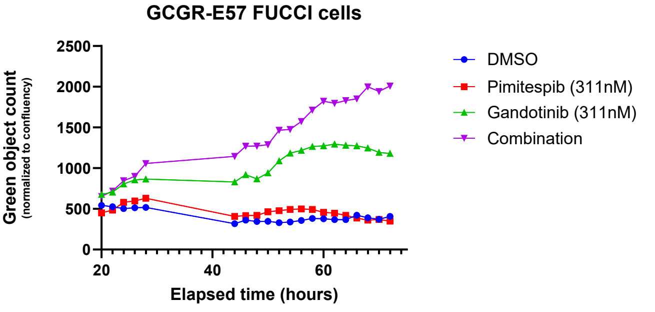


**Supplementary Figure 5. Combined JAK2 and HSP90 inhibition alters cell cycle progression in GCGR-E57 cells.**
Left: representative live-cell images of GCGR-E57 FUCCI cells treated with DMSO, Gandotinib, Pimitespib, or the combination for 72 hours. Right: quantification of green FUCCI-positive object count normalised to cell confluency over time.

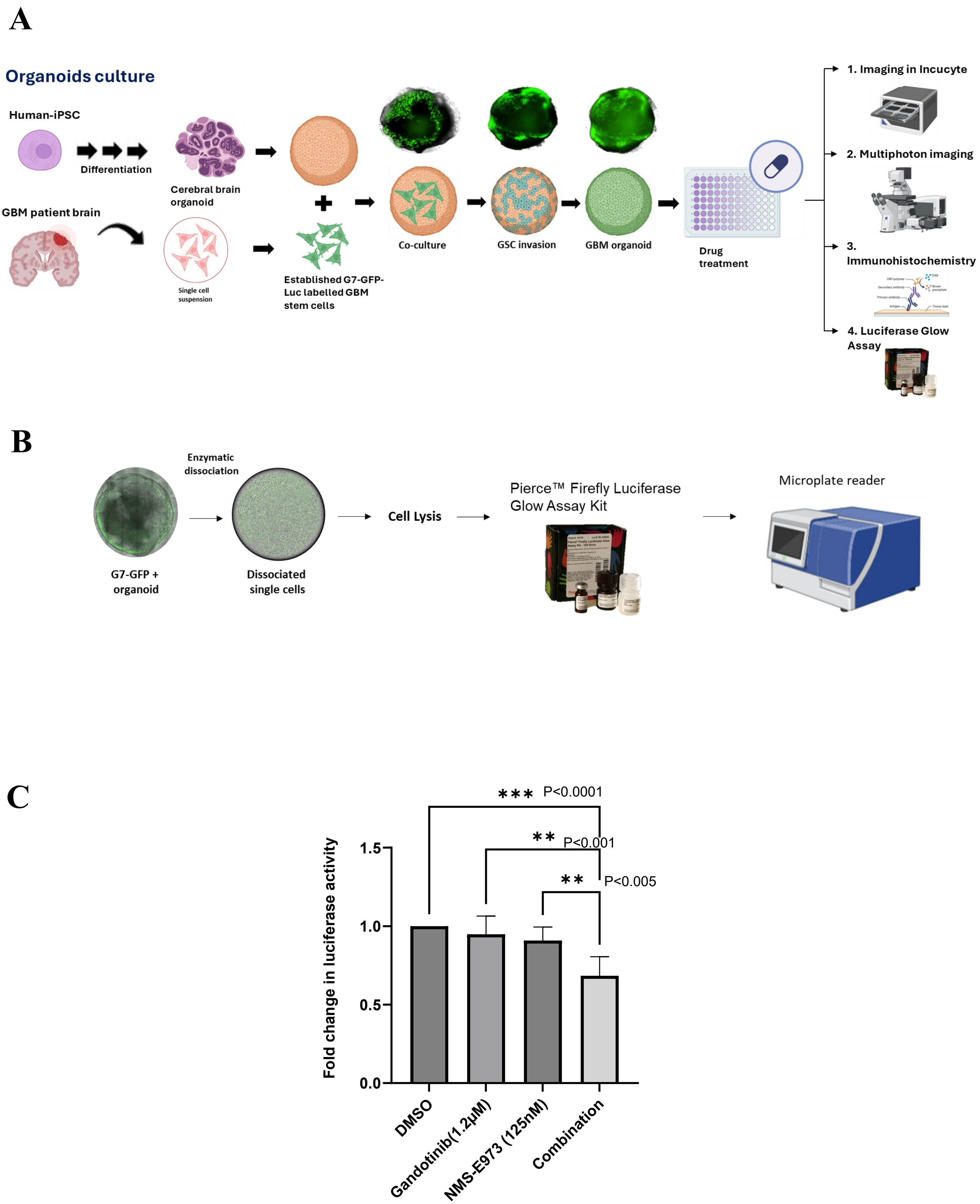


**Supplementary Figure 6. Luciferase-based assessment of tumour burden in GBM–cerebral organoid co-cultures.** **(A)** Schematic showing the generation of human iPSC-derived cerebral organoids, co-culture with established patient-derived G7 GBM stem-cell line expressing GFP-Luc followed by therapeutic treatment, and multimodal assessment by live-cell imaging, multiphoton microscopy, histology and luciferase assay. **(B)** Schematic overview of luciferase based tumour growth assay. G7-GFP-Luc organoid co-cultures were enzymatically dissociated, lysed, and analysed using a firefly luciferase glow assay to quantify viable luciferase-labelled tumour cells according to the manufacturer's instructions. **(C)** Quantification of tumour burden by luciferase activity following 2 weeks of treatment. Data are presented as fold change in luciferase activity relative to the DMSO control (n = 5 organoids per technical replicate of three independent experiments). Statistical significance was assessed using one-way ANOVA followed by Dunnett's multiple comparisons test. Adjusted *P* values are indicated on the graph. Schematics in (A) and (B) were created in BioRender. Elliott, R. (2027) https://BioRender.com/tqqlqhe.

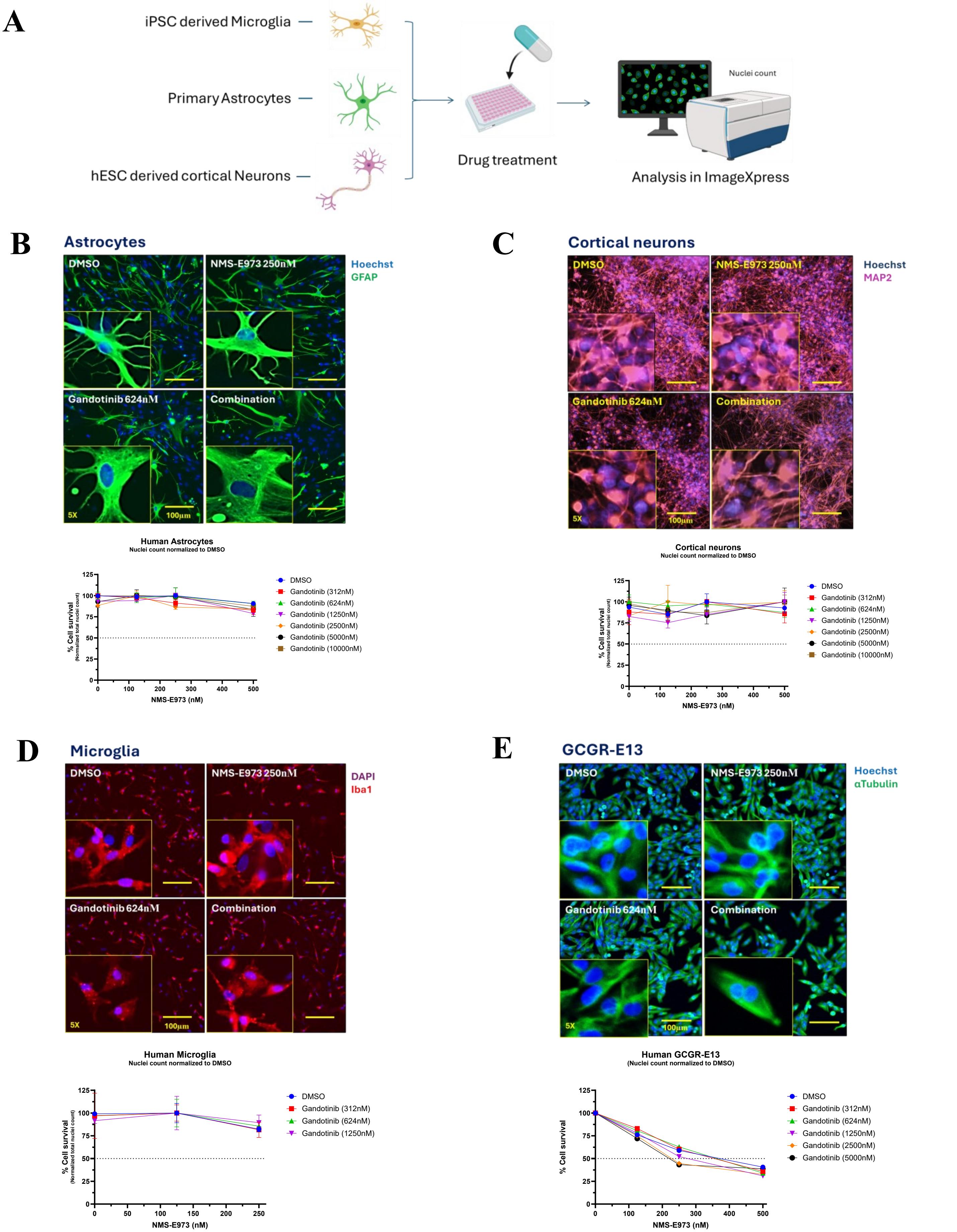


**Supplementary Figure 7. JAK2 and HSP90 inhibitor combinations show lower sensitivity in primary human astrocytes, iPSC-derived microglia and hESC-derived cortical neurons.**
**(A)** Schematic of neural cell-type drug-response analysis. **(B–E)** Representative immunofluorescence images and corresponding dose-response curves for **(B)** primary human astrocytes, **(C)** hESC-derived cortical neurons, **(D)** iPSC-derived microglia and **(E)** GCGR-E13 GBM cells treated with NMS-E973 in combination with increasing concentrations of Gandotinib for 72 hours. Astrocytes were stained for GFAP, cortical neurons for MAP2 and microglia for Iba1; nuclei were stained with Hoechst. Schematic in (A) was created in BioRender. Elliott, R. (2027) https://BioRender.com/ryvc2gg.

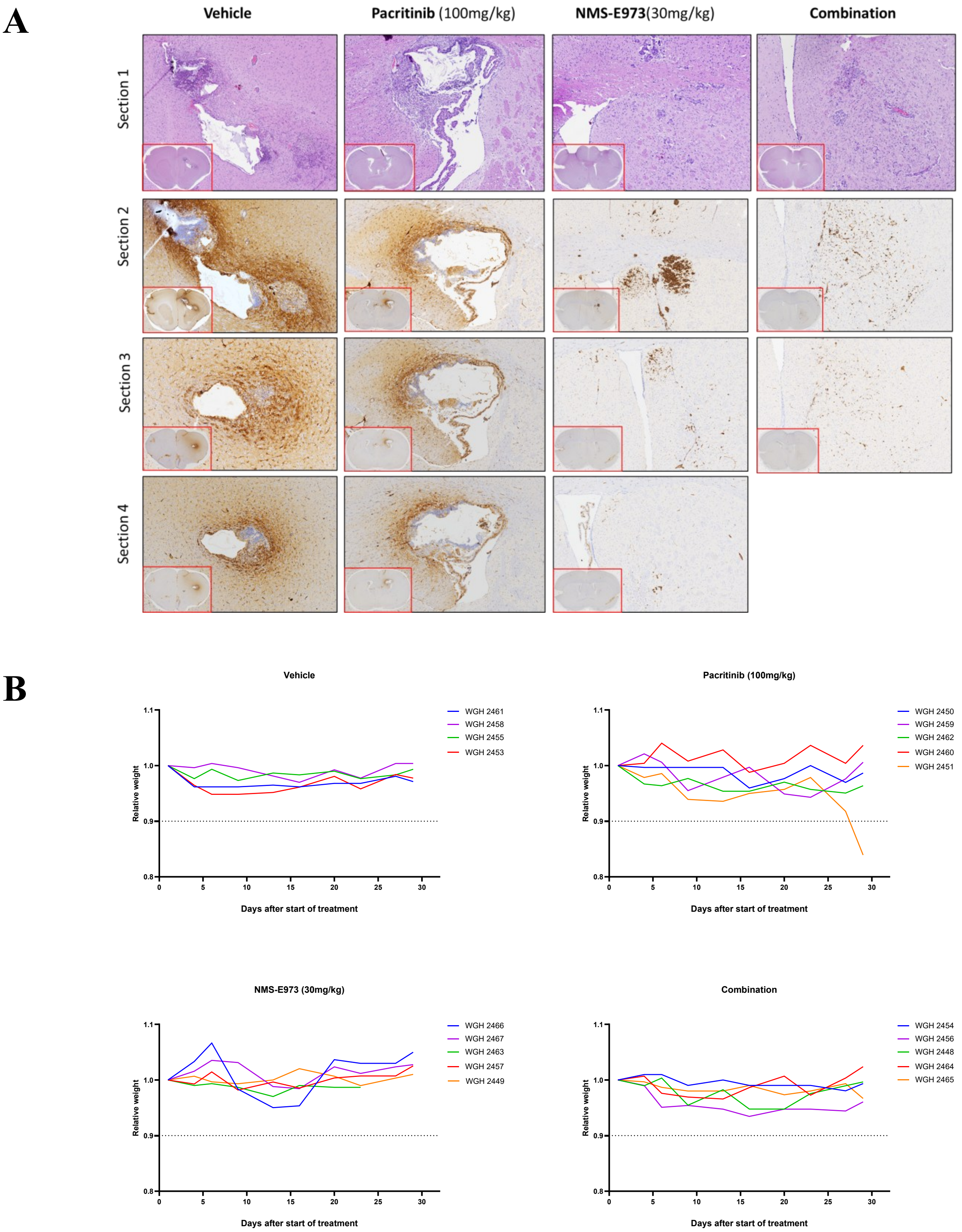


**Supplementary Figure 8. Histological assessment of tumour burden and body-weight monitoring in G7 intracranial xenografts. (A)** Representative H&E and human mitochondria-immunostained brain sections (4 μm) from individual mice treated with vehicle, Pacritinib (100 mg/kg), NMS-E973 (30 mg/kg), or the combination. Multiple adjacent sections from each brain are shown. **(B)** Relative body weight of individual mice over the treatment period.

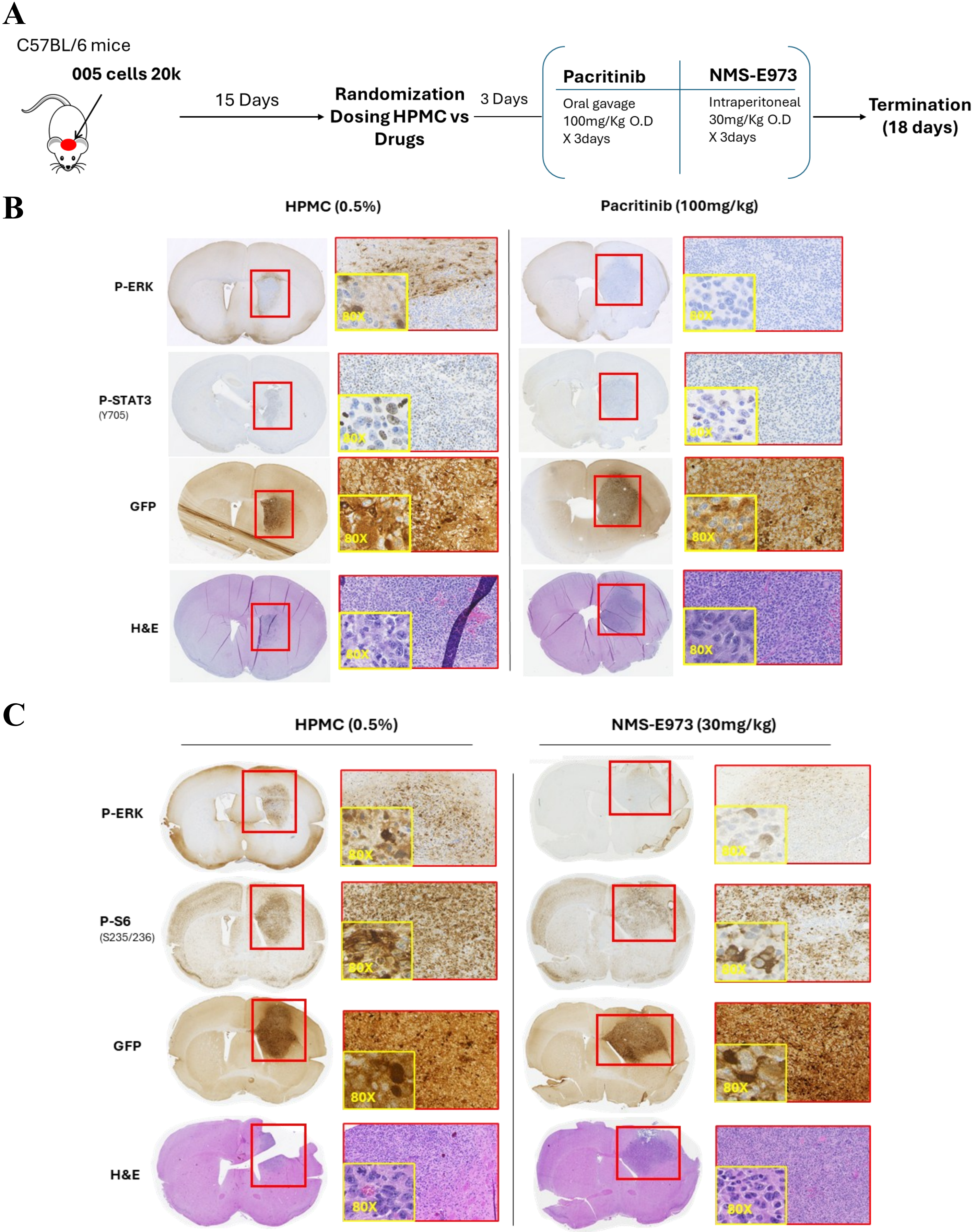


**Supplementary Figure 9. Pharmacodynamic analysis of Pacritinib and NMS-E973 in 005-derived intracranial tumours.** **(A)** Schematic of the *in vivo* pharmacodynamic study. GFP-labelled 005 cells were implanted intracranially into C57BL/6 mice. Mice were randomised 15 days after implantation and treated with vehicle or drug for 3 days before brain collection at day 18. **(B-C)** Representative H&E and immunohistochemistry-stained brain sections from vehicle **(B)** Pacritinib and **(C)** NMS-E973 treated mice. Pacritinib treated brain sections were stained for p-ERK and p-STAT3 Y705 while NMS-E973 stained for p-ERK and p-S6 Ser235/236 to assess the pathway modulation. GFP staining identifies tumour bearing regions in all mice.
